# A truncated reverse transcriptase enhances prime editing by split AAV vectors

**DOI:** 10.1101/2021.11.05.467423

**Authors:** Zongliang Gao, Jakob Haldrup, Sujan Ravendran, Nanna S. Mikkelsen, Jacob Giehm Mikkelsen, Rasmus O. Bak

## Abstract

Prime editing is a new CRISPR-based genome editing technology that relies on the prime editor (PE), a fusion protein of Cas9-nickase and M-MLV reverse transcriptase (RT), and a prime editing guide RNA (pegRNA) that serves both to target PE to the desired genomic locus and to carry the edit to be introduced. Here, we make advancements to the RT moiety to improve prime editing efficiencies and truncations to mitigate issues with AAV viral vector size limitations, which currently do not support efficient delivery of the large prime editing components. These efforts include RT variant screening, codon optimization, and PE truncation by removal of the RNase H domain and further trimming. This led to a codon-optimized and size-minimized PE that has an expression advantage (1.4× fold) and size advantage (621 bp shorter). In addition, we optimize the split intein PE system and identify Rma-based Cas9 split sites (573-574 and 673-674) that combined with the truncated PE delivered by dual AAVs result in superior AAV titer and prime editing efficiency. This novel minimized PE provides great value to AAV-based delivery applications *in vivo*.

## Introduction

Prime editing enables versatile and precise genome editing independent of double-strand DNA breaks and exogenous donor template DNA and thus exhibits great potential as a tool for biomedical research and gene therapy^1^. The most efficient prime editing system consists of: (i) a prime editor (PE), an SpCas9 nickase (nCas9) fused to a mutant Moloney Murine Leukemia Virus (M-MLV) reverse transcriptase (RT) (**Fig. 1A**), (ii) a prime editing guide RNA (pegRNA) which carries the desired edit, and (iii) a nicking guide RNA (ngRNA). Despite of its versatile genome editing capacity, prime editing is challenged by the large PE gene (~6.4 kb), which makes delivery of the prime editing system (PE, pegRNA, and ngRNA) challenging. For instance, it is impossible to package the prime editing system into a single adeno-associated virus (AAV) vector, which is limited to ~5 kb with the inclusion of the inverted terminal repeats (ITRs). Although two recent studies reported that the PE gene can be divided in two and delivered by dual AAV vectors relying on trans-splicing intein^2, 3^, the large size of PE severely restricts the position of the split site, which leads to suboptimal prime editing efficiency and leaves limited space for additional functional elements.

**Figure 1.**
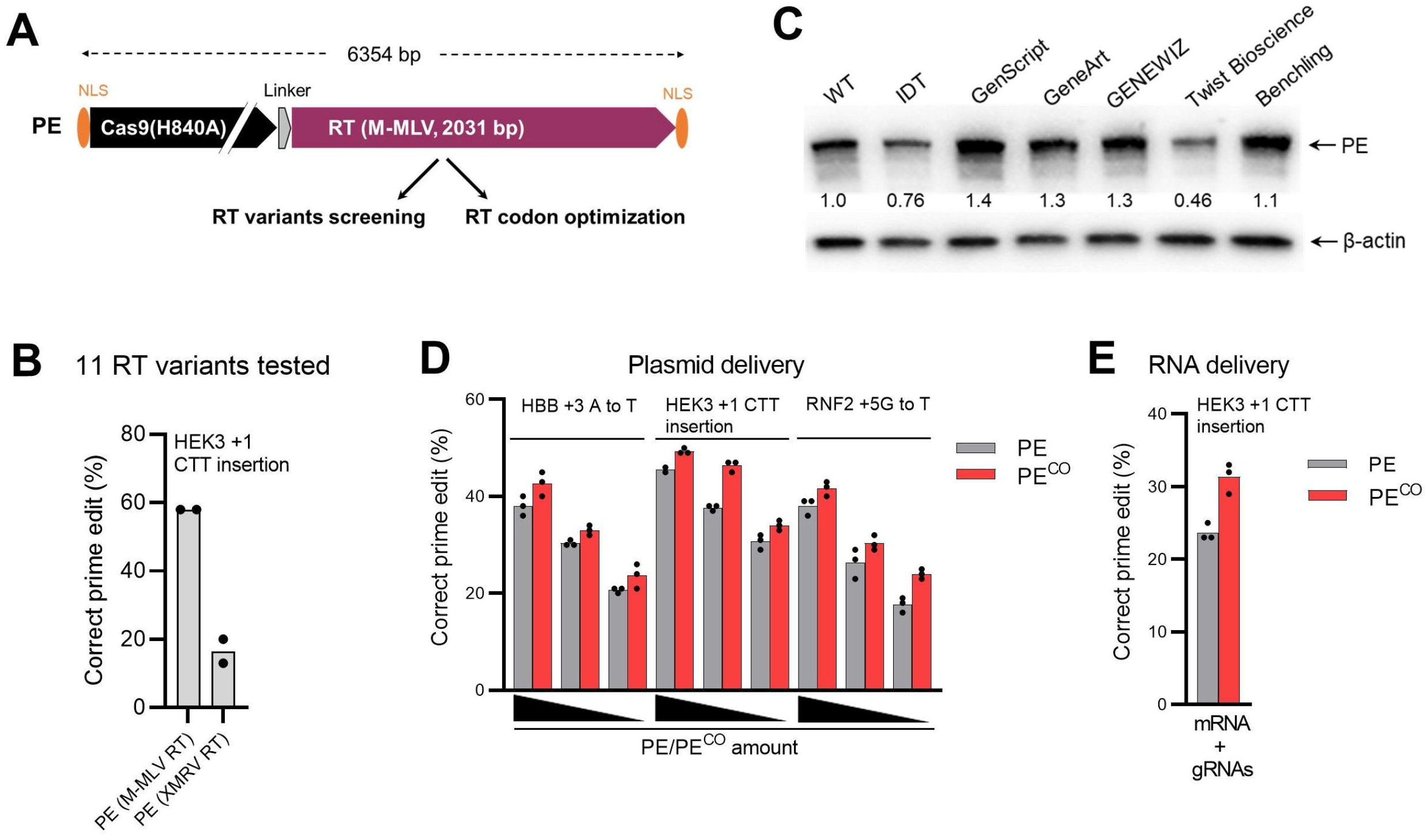
RT variant screening and codon optimization. (**A)** Schematic of the PE gene cassette employed in this study. PE consists of Cas9-nickase (Cas9 (H840A)) and M-MLV RT. The bipartite SV40 nuclear localization signals (bpNLS) are marked in orange and a linker in gray. (**B**) Screening of prime editing activity of eleven RT variants codon-optimized by GenScript showed activity only for XMRV RT (all variant data shown in **Fig. S1**). PE constructs were transfected with plasmids encoding pegRNA and ngRNA into HEK293T cells and prime editing results were analyzed after 3 days. **(C)** The effect of RT codon optimization on PE expression using algorithms from different companies. Same amounts of PE plasmids were transfected into HEK293T cells and western blotting was conducted after 3 days. PE protein expression levels were normalized to β-actin protein levels and wild type (WT) PE protein expression level was arbitrarily set to 1.0 (indicated below the PE blot). The image shows a representative blot from two independent experiments with similar results. (**D**) Comparison of prime editing frequencies of the original PE and the codon-optimized PE^CO^ by plasmid delivery. HEK293T cells were transfected with varying amounts of PE or PE^CO^ (1500, 300, and 60 ng) together with fixed amounts of plasmid encoding pegRNA and ngRNA. After 3 days, PCR products were subjected to Sanger sequencing and ICE analysis to evaluate prime editing frequencies. **(E)** Comparison of prime editing frequencies of PE and PE^CO^ by all-RNA delivery. PE mRNA and synthetic pegRNA and ngRNA were electroporated into HEK293T cells. Cells were subjected to ICE analysis 3 days post-transfection. The data in the graph are presented by mean values with all data points from independent experiments shown.

In addition, the efficiency of prime editing can vary greatly across edit types, target loci, and cell types. Hence, there is a need for an improved prime editing system. Very recently, prime editing activity improvements were achieved by pegRNA engineering, nuclear localization signal (NLS) optimization to the PE, and by manipulation of cellular DNA repair pathways^4, 5, 6^.

In this study, by engineering the RT of the PE, we developed a codon-optimized and size-minimized PE. Combining the optimized intein split PE system, a dual AAV split PE system was generated that has vastly improved AAV titer and prime editing activity.

## Results

### M-MLV RT performs the best among RT variants

With an attempt to improve prime editing activity and obtain a shorter PE with a RT of reduced size, we tested if RTs from other species could replace M-MLV RT to constitute a functional PE. A screening of 11 human codon-optimized RT variants, including 9 retroviral RTs and 2 bacterial group II intron RTs (Supplementary Data 1), showed that only Xenotropic murine leukemia virus-related virus (XMRV) RT is functional in inducing prime editing (HEK3 +3 CTT insertion) in the HEK3 gene (**Fig. 1B and Fig. S1**). However, this XMRV RT-containing PE induced much lower prime editing than that of M-MLV RT (17% vs. 58%).

### RT codon optimization improves prime editing activity

We next attempted to improve prime editing activity by increasing PE expression level. Since the SpCas9 has been widely developed, we focused on the RT moiety. We generated M-MLV RT variants using 6 different codon optimization algorithms from different companies: IDT, GenScript, GeneArt, GeneWIZ, Twist Bioscience, and Benchling. 4 out of 6 were observed to increase PE protein expression level compared to the previously published variant with GenScript (hereafter termed PE^CO^) expressing highest protein levels (1.4-fold increase) (**Fig. 1C**). We next investigated the performance of PE^CO^ by plasmid delivery of varying PE and PE^CO^ amounts to HEK293T cells. Results consistently showed that PE^CO^ exhibited higher prime editing frequencies at all three tested targets (HBB, HEK3, and RNF2) (**Fig.1D**). We also performed this comparison with all-RNA delivery (PE mRNA with chemically modified pegRNAs and ngRNA), which is more therapeutically relevant in primary cells, and we observed a similar increase in editing by PE^CO^ (31.3% vs. 23.6%) (Fig. 1E).

### Engineering a minimal M-MLV RT with uncompromised prime editing activity

Next, we sought to shorten the size of M-MLV RT. RT consists of an RNA-dependent polymerase domain and an RNase H domain (Fig. 1A). The polymerase domain uses single strand RNA as template to synthesize single strand DNA, and the RNase H domain selectively degrades the RNA template of RNA/DNA hybrids. Considering that the RNase H domain is only for RNA template degradation and that this domain is in fact inactive in the mutant M-MLV RT of PE, we hypothesized that the RNase H domain would be dispensable. We also hypothesized that the polymerase region might be trimmed at both ends while still retaining full activity. To test this, we deleted the RNase H domain of M-MLV RT (471 bp) to make a truncated PE^CO^ (PE^CO^-ΔR) and based on PE^CO^-ΔR we made further truncations by 30 bp serial N- or C-terminal truncations to RT, creating a total of eleven different truncated PE^CO^ variants (**Fig. 2A**). Prime editing experiments in HEK293T cells for two targets (HEK3 and HBB) showed that PE^CO^-ΔR variants with a maximally 60 bp N-terminal deletion and 120 bp C-terminal deletion did not affect prime editing activity compared to the PE^CO^-ΔR variant. (**Fig. 2B**). We then tested four combinatorial N- and C-terminal truncations and showed that a combined 150 bp deletion (ΔN60+C90) to PE^CO^-ΔR did not affect the prime editing efficiency compared to the PE^CO^-ΔR variant, leading to a minimal PE (PE^CO^-Mini) (**Fig. 2C**). To finally assess the performance of PE^CO^-Mini we performed a parallel comparison to the full-length PE^CO^ and PE^CO^-ΔR. These data showed that the PE^CO^-Mini displayed very similar prime editing frequencies at three different target genes (**Fig. 1D**). Furthermore, we observed no difference in the editing outcome (specific prime edits and unspecific indels) of the three PE variants (**Fig. S2**). These results show that the RNase H domain is dispensable to PE, which was further supported by the observed functionality of an RNase H-deleted XMRV RT PE variant (**Fig. S1**). Intriguingly, PE with XMRV RT-ΔR outperformed XMRV RT (functional RNase H domain) in prime editing in the HBB gene (17% vs. 28%), indicating a negative effect from an active RNase H domain. One could speculate that this domain might degrade the pegRNA region where it anneals to the RT primer binding site (PBS) of the DNA target. However, we still did not observe any prime editing from other RNase H-deleted RT PE variants (**Fig. S1**). In conclusion, we engineered a codon-optimized and size-minimized PE^CO^-Mini with an overall size reduction of 621 bp that retains prime editing activity compared to the full-length PE.

**Figure 2.**
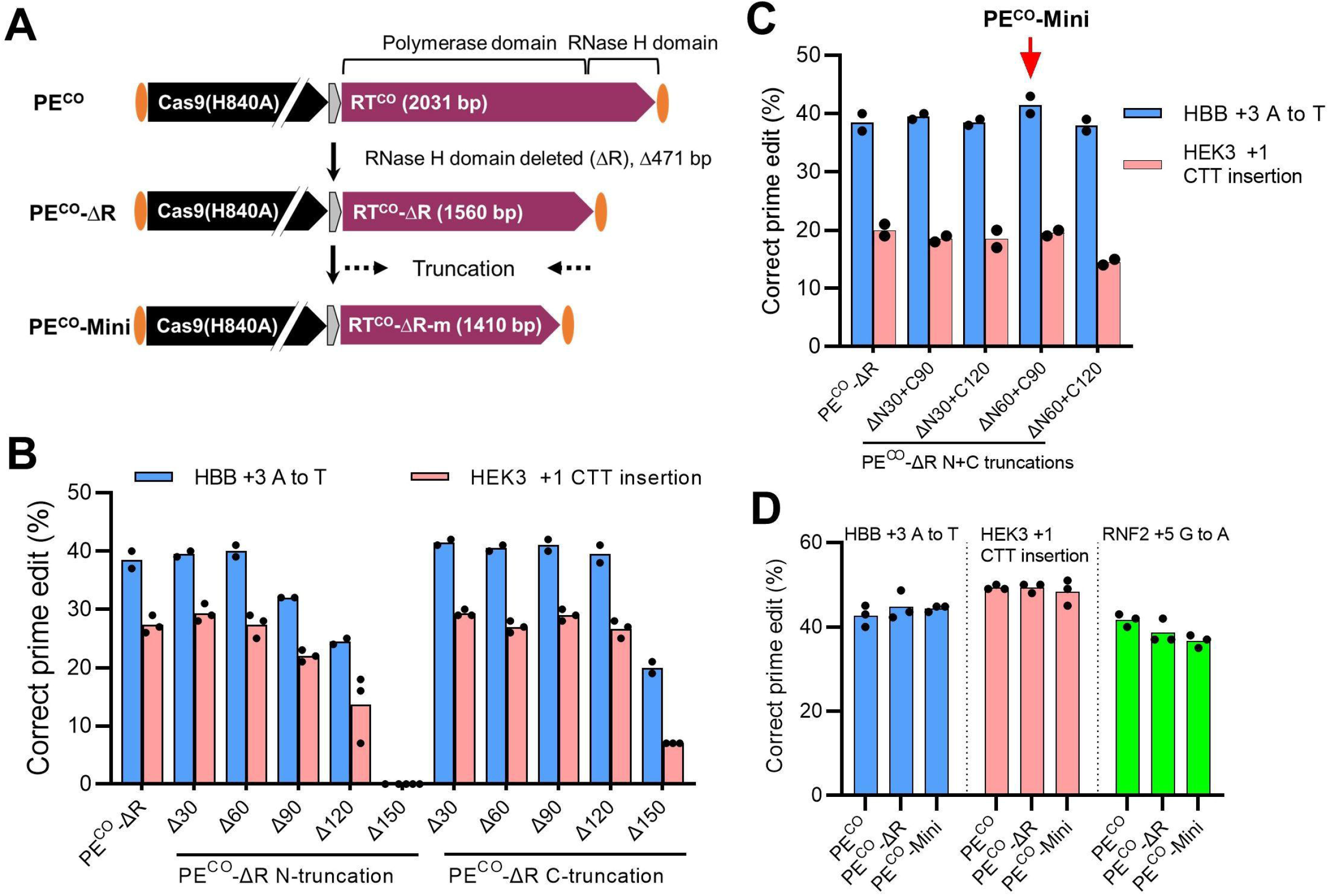
Generation of a minimal PE by truncating M-MLV RT. (**A**) Schematic of PE^CO^ and truncated PE^CO^ variants used in this study. The RT is composed of a polymerase and an RNase H domain. Initially, a truncated PE was made with the RNase H domain deleted (471 bp) to form PE^CO^-ΔR. Subsequently, serial 30 bp truncations at both ends were tested with the aim of generating a minimal PE^CO^ (PE^CO^-Mini). (**B**) Minimizing M-MLV RT by deleting the RNase H domain (PE^CO^-ΔR) and by further N- and C-terminal trimming. The same molar amounts of PE^CO^-ΔR and trimmed PE^CO^-ΔR plasmids were transfected into HEK293T cells together with fixed amounts of plasmids encoding pegRNA and ngRNA. Cells were subjected to ICE analysis for evaluating prime editing activity 3 days post-transfection. (**C**) Combinatorial N- and C-terminal truncations were analyzed for prime editing efficiencies as in (**B**). Based on the activity, we selected RT-ΔR with ΔN60+C90 as the minimal, which is 1410 bp long. (**D**) Comparison of PE^CO^ and the two truncated variants PE^CO^-ΔR and PE^CO^-Mini performed as in (**B**). The data are presented by mean values with all data points from independent experiments shown.

### Optimization of an intein split PE system

To pave the way for effective dual AAV delivery of PE, it is important to establish an efficient split PE system. To do this, we constructed various intein-based split PE^CO^ systems (**Fig. 3A**). Previous studies have shown that Cas9 activity is highly dependent on the position of the intein split site. We therefore chose four commonly used Cas9 split sites (573-574, 637-638, 674-675, and 713-714) and two different inteins (Npu and Rma)^7, 8^. The split and non-split PE^CO^ systems were tested at the HBB and HEK3 targets in HEK293T cells. The efficiency of the eight two-plasmid split PE^CO^ systems varied markedly, and as expected all were lower than the single PE^CO^ plasmid (no splitting) (**Fig. 3B**). Rma 573-574 was the most efficient (85% activity of that of single-plasmid full-length PE^CO^) followed by Rma 674-675 (75% activity) (**Fig. 3B**). An AAV genome size calculation showed that if using the two most efficient intein sites (Rma 573-574 and 674-675) to split non-truncated PE^CO^, the PE^CO^ C-terminal part would amount to >5 kb thereby exceeding the ~5 kb AAV packaging limit (including ITRs)^9^, even with the use of a small promoter (EFS) and poly(A) signal (**Fig. 3A**). However, the two intein split sites could be used with the PE^CO^-Mini with vector genome sizes below 4.7 kb. Despite the slightly higher activity of the Rma 573-574 split site, we chose the Rma 674-675 split site as it leaves 300 bp of space in the vector encoding the C-terminal part, which would allow a more flexible promoter choice (total AAV vector size of 4.4 kb) (**Fig. 3C**). Further analysis of plasmid-based delivery of split PE^CO^, PE^CO^-ΔR, and PE^CO^-Mini systems with the Rma 674-675 split site showed similar editing activity for all three targets tested, which is around 75% of the single all-in-one PE^CO^ plasmid system (**Fig. 3D**).

**Figure 3.**
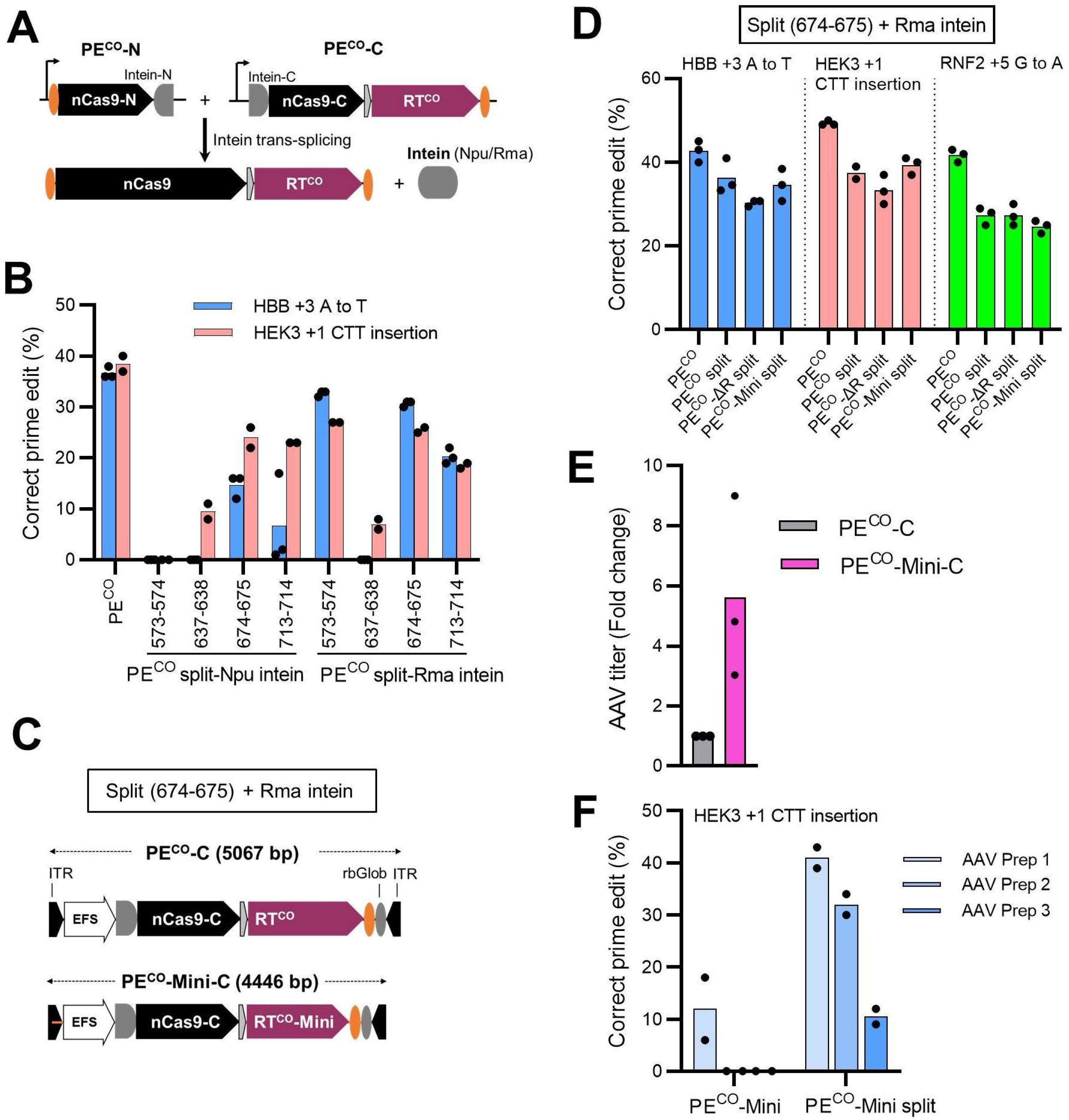
The engineered minimal PE variant enhances prime editing when delivered by dual AAV vectors. **(A)** Schematic of the split PE system where the full-length PE is reconstituted by intein protein-mediated trans-splicing. **(B)** Screening of split PE systems with different inteins and Cas9 split positions by plasmid transfection of HEK293T cells. **(C)** Schematic of AAV vector genomes encoding the C-terminal part of the split PEs (PE^CO^-C or PE^CO^-Mini-C). ITR: inverted terminal repeats; rbGlob: rabbit beta globin poly(A) signal. **(D)** Comparable activity of the split full-length and the two split truncated PE systems (PE^CO^-ΔR split and PE^CO^-Mini split) by plasmid transfection in HEK293T cells. The single plasmid non-truncated PE^CO^ was included for comparison. **(E)** AAV6 vectors encoding the C-terminal part of the split PE^CO^-Mini (PE^CO^-Mini-C) were produced in parallel with vectors encoding the C-terminal part of the non-truncated PE^CO^ (PE^CO^-C). AAV titers were determined by ddPCR with the titers of PE^CO^-C set to 1. **(F)** HEK293T cells were transduced with equal MOIs (2×10^5^ vg/cell) of the generated split AAV vectors and prime editing frequencies were evaluated five days post-transduction by Sanger sequencing and ICE analysis. The data are presented by mean values with all data points from independent experiments shown.

### The truncated PE enhances PE system delivery by AAV vectors

To investigate the performance of the PE^CO^-Mini compared to full-length PE^CO^ for AAV delivery, we packaged dual AAV6 vectors based on the Rma 674-675 split site (**Fig. 3C and Fig. S3**) and measured the vector titers and performed prime editing assays. RNA Pol III cassettes for the pegRNA (U6 promoter) and ngRNA (7SK promoter) were placed in the vector encoding the PE N-terminal part amounting to a vector genome size of 3.5 kb (**Fig. S3**). AAV vectors were produced in parallel and titer measurements by ddPCR showed a >5-fold higher titer of the vector carrying the C-terminal part of PE^CO^-Mini (PE^CO^-Mini-C; 4.4 kb) compared to the vector carrying the C-terminal part of the full-length PE^CO^ (PE^CO^-C; 5.1 kb) (**Fig. 3E**). Transduction of HEK293T cells using the same MOI of split PE^CO^ and split PE^CO^-Mini systems showed 10-40% editing with the split PE^CO^-Mini system compared to 0-10% editing by the split PE^CO^ system (**Fig. 3F**). Since cells were transduced at equal MOIs, this indicates that the PE^CO^-C vector genomes are too large to produce functional vector particles, which leads to inefficient PE^CO^ expression in target cells. Thus, PE^CO^-Mini is highly advantageous for dual AAV split PE delivery.

## Discussion

The versatile gene editing ability of prime editing not only makes it a very useful biomedical research tool, but also holds great promise as a therapeutic approach to correct genetic mutations. However, the prime editing efficiency greatly varies across target loci and cell types, impeding its broad application^1^. Importantly, the large size of PE makes delivery very challenging, especially for *in vivo* application where AAV vectors constitute a popular vector choice but suffer from limited packaging capacity. Although PE system delivery has been achieved by intein-mediated splicing between dual AAV vectors carrying a split PE, the large size of the PE and the AAV packaging limitation severely restrict the position where the intein can be placed in the PE gene. Consequently, this leads to suboptimal prime editing efficiency and leaves limited space for additional functional elements in the vectors.

In this study, we engineered the M-MLV RT to develop a codon-optimized and minimal PE: PE^CO^-Mini. Screening of several RT variants from other species revealed only XMRV RT to be functional but displaying much less activity than the M-MLV. Codon-optimization to RT increased PE expression level by 1.4-fold and consequently improved prime editing activity as demonstrated by both plasmid and all-RNA delivery. Removal of the RNase H domain and subsequent trimming of RT resulted in a minimal PE that is 621 bp shorter than the original PE but maintains editing efficiency. By testing different Cas9 split sites and inteins, we identified two split sites (573-574 and 673-674) together with the Rma intein to be the most potent split PE systems. We demonstrated that when using 673-674/Rma, PE^CO^-Mini enables a split PE system for AAV packaging in which the two vector genomes are well below the AAV packaging limit in contrast to non-truncated PE^CO^ (**Fig. 3D**). As a result, PE^CO^-Mini leads to superior AAV titers and prime editing activity following PE delivery by AAV transduction.

Intein-mediated split PE system delivery by dual AAV vectors enables versatile gene editing for in vivo gene therapy. We show here that choice of intein and the position of the Cas9 split site is critical to prime editing efficiency, similar to the findings for base editing^7, 8^. For instance, a previous publication employed the Npu intein and Cas9 713-714 split site for PE delivery by dual AAV vectors^2^, but in our hands this system displayed suboptimal prime editing efficiency (**Fig. 3B**). Although we have identified two efficient split positions for PE delivery, a larger screening of Cas9 split sites and inteins might identify a more robust split PE system. Such investigations can be reinforced by the extra 621 bp space provided by the PE^CO^-Mini. This extra capacity allows the accommodation of a more flexible promoter choice, e.g. the use of tissue-specific promoters or other functional elements. Similar to the beneficial effect to AAV vectors, we believe that PE^CO^-Mini would also benefit delivery by other viral vector systems like lentiviral vectors where the reduced vector size is known to improve vector titers^10^.

## Acknowledgements

ZG gratefully acknowledges support from an individual postdoctoral fellowship from the Lundbeck Foundation (R303-2018-3571). ROB gratefully acknowledges the support from a Lundbeck Foundation Fellowship (R238-2016-3349), the Independent Research Fund Denmark (0134-00113B, 0242-00009B, and 9144-00001B), an AIAS-COFUND (Marie Curie) fellowship from Aarhus Institute of Advanced Studies (AIAS) co-funded by Aarhus University’s Research Foundation and the European Union’s seventh Framework Program under grant agreement no 609033, the Novo Nordisk Foundation (NNF19OC0058238 and NNF17OC0028894), Innovation Fund Denmark (8056-00010B), the Carlsberg Foundation (CF20-0424 and CF17-0129), Slagtermester Max Wørzner og Hustru Inger Wørzners Mindelegat, the AP Møller Foundation, the Riisfort Foundation, and a Genome Engineer Innovation Grant from Synthego.

## Author contributions

ZG and ROB conceived the study and designed the experiments. ZG performed the experiments and analyzed the data with assistance from JH, SR, and NSM. JGM and ROB supervised the study. ZG and ROB wrote the manuscript. All authors reviewed, edited, and approved the final manuscript.

## REFERENCES

1. Anzalone AV, et al. Search-and-replace genome editing without double-strand breaks or donor DNA. Nature 576, 149–157 (2019).

2. Liu P, et al. Improved prime editors enable pathogenic allele correction and cancer modelling in adult mice. Nat Commun 12, 2121 (2021).

3. Zhi S, et al. Dual-AAV delivering split prime editor system for in vivo genome editing. Molecular Therapy, (2021).

4. Liu Y, et al. Enhancing prime editing by Csy4-mediated processing of pegRNA. Cell Res 31, 1134–1136 (2021).

5. Nelson JW, et al. Engineered pegRNAs improve prime editing efficiency. Nat Biotechnol, (2021).

6. Chen PJ, et al. Enhanced prime editing systems by manipulating cellular determinants of editing outcomes. Cell, (2021).

7. Chen Y, et al. Development of Highly Efficient Dual‐AAV Split Adenosine Base Editor for In Vivo Gene Therapy. Small Methods 4, 2000309 (2020).

8. Levy JM, et al. Cytosine and adenine base editing of the brain, liver, retina, heart and skeletal muscle of mice via adeno-associated viruses. Nature biomedical engineering 4, 97–110 (2020).

9. Wu Z, Yang H, Colosi P. Effect of genome size on AAV vector packaging. Mol Ther 18, 80–86 (2010).

10. Gao Z, Herrera-Carrillo E, Berkhout B. A single H1 promoter can drive both guide RNA and endonuclease expression in the CRISPR-Cas9 system. Molecular Therapy-Nucleic Acids 14, 32–40 (2019).

