## Supplemental information for "A truncated reverse transcriptase enhances prime editing by split AAV vectors"

**SUPPLEMENTARY INFORMATION**

**Supplementary Figures**

### Figure S1

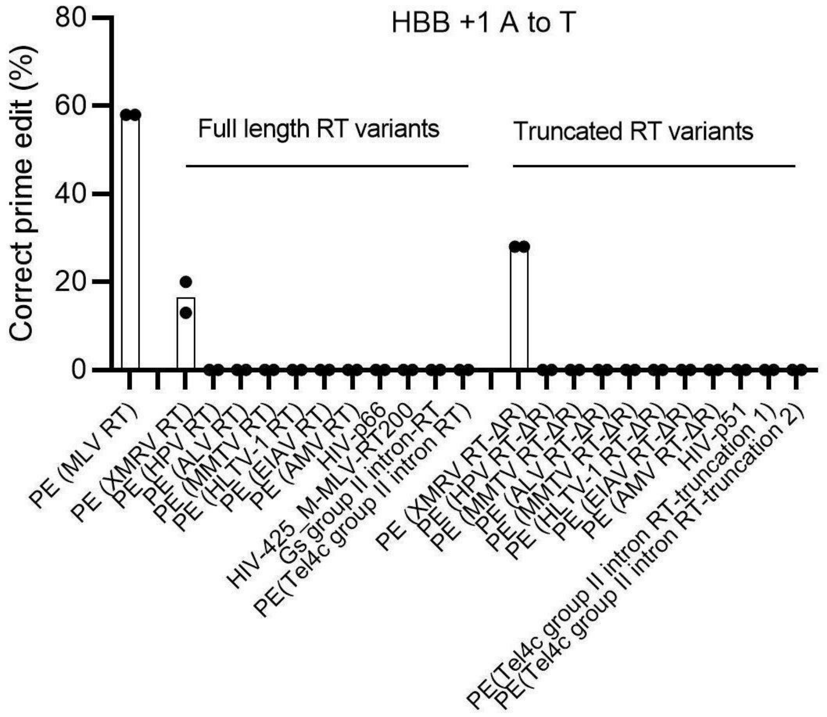

### Figure S2

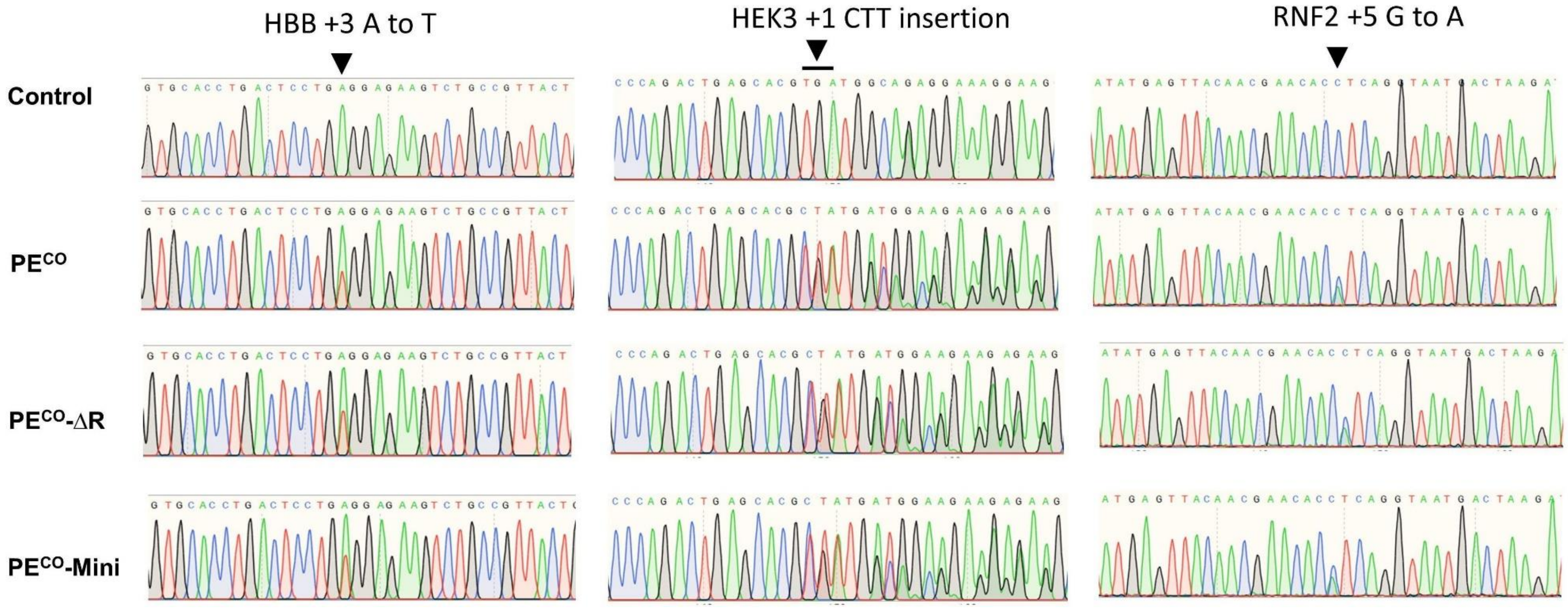

HBB +3 A to T
HEK3 +1 CTT insertion
RNF2 +5 G to A
Control
PE^CO
PE^CO-ΔR
PE^CO-Mini

**Figure S3**

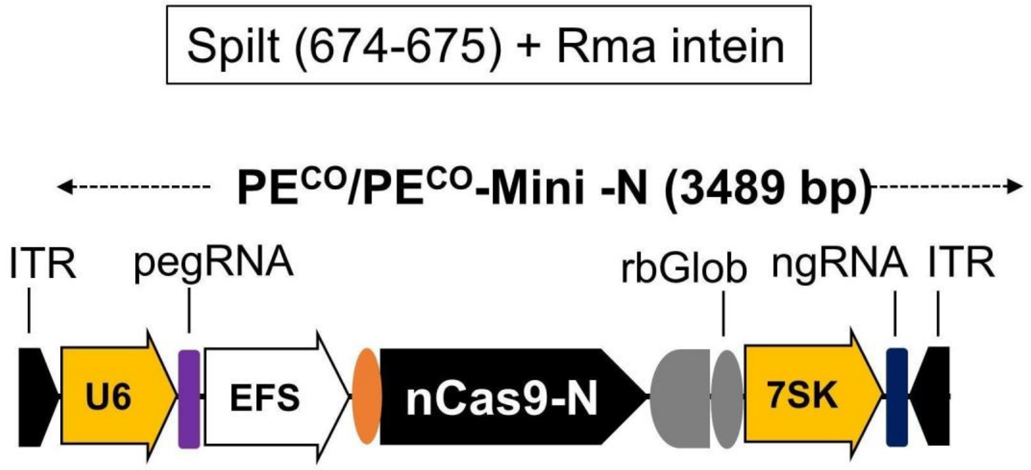

Spilt (674-675) + Rma intein
PECO/PECO-Mini -N (3489 bp)
ITR
pegRNA
rbGlob
ngRNA
ITR
U6
EFS
nCas9-N
7SK

#### Materials and Methods

##### Plasmid construction

The plasmids pCMV-PE2 (addgene #132775) expressing prime editor 2 and pU6-pegRNA-GG-acceptor (addgene #132777) expressing pegRNA were gifts from David Liu.[1] To test the RT variants from other species, the 12 selected RT variants were human codon-optimized by GenScript and then the corresponding DNA fragments were ordered from Twist Bioscience, which were inserted into the PE2 plasmid to replace M-MLV RT by Gibson cloning using BamHI and AgeI restriction enzyme sites. To make PE constructs with RTs of different codon usage, M-MLV RT was codon-optimized by different web sources and the DNA fragments were ordered from Twist Biosciences and inserted into PE2 by BamHI and AgeI sites. The DNA sequences of RT variants can be found in Supplemental Data 1 and 2. The DNA sequences of two truncated M-MLV RT are listed in Supplemental Data 3. pegRNA plasmids were cloned following the protocol from David Liu.[1] Nicking gRNA (ngRNA) expression plasmid was cloned using a home-made px330 plasmid (addgene plasmid #42230; a kind gift from Feng Zhang) in which Cas9 was removed and the sgRNA expressed from the U6 promoter. Nucleotides encoding ngRNA and pegRNA spacers and 3' extension sequences are listed in Supplementary Table 1.

AAV vector plasmids were cloned pAAV-MCS (Agilent technologies) containing inverted terminal repeats from AAV2. The AAV plasmids were created by Gibson cloning, of which the DNA fragments encoding the split PE system were PCR-amplified or synthesized by Twist Bioscience. The schematic of AAV genome are displayed in **Fig. S3**.

##### IVT (In Vitro Transcription)

IVT of PE mRNA was performed as described previously.[2] Briefly, plasmids for IVT were made based on a backbone containing the T7 promoter, gene of interest (PE), a murine Hba-a1 3'UTR, and a stretch of 50 adenines [poly(A)]. The plasmid was then linearized with a BtgZI restriction enzyme cutting immediately after the poly(A). The linearized plasmid was concentrated and purified by Ammonium Acetate precipitation. IVT reactions were conducted using the MEGAscript Kit (Ambion, Thermo Fisher Scientific) according to the manufacturer's instructions, but with full substitution of uridine with pseudouridine (Trilink Biotechnologies or APExBio) and co-transcriptional capping with CleanCap AG (Trilink Biotechnologies) in a 1:4 ratio between GTP and CleanCap. The mRNA was purified and concentrated using the RNA Clean and Concentrator kit (Zymo Research). The quality of the IVT RNA was confirmed on a denaturing formaldehyde gel and quantified by UV-Vis spectrophotometry.

**sgRNAs**

Synthetic pegRNA and ngRNA were purchased from Synthego as chemically modified sgRNAs containing 2'-O-Methyl groups at the three first and last bases and 3' phosphorothioate bonds between the first 3 and the last 2 bases.

**Cell culture**

HEK293T cells were cultured in DMEM (Life Technologies) supplemented with 10% fetal calf serum (FCS), 2 mM L-glutamine, penicillin (100 U/mL), and streptomycin (100mg/mL).

**Transfection / Electroporation**

Plasmid transfection was conducted by Lipofectamine 2000 (Invitrogen). Briefly, 0.5 ml culture medium with 1.3x $10^5$ cells were seeded per well in 24-well plates one day prior to transfection. Transfections were performed with Lipofectamine 2000 according to the manufacturer's protocol. Unless specified, 1500 ng PE plasmid, 500 ng pegRNA plasmid, and 245 ng ngRNA were used per well. For comparing the split and non-split PE systems, a same molar of split or non-split PE plasmids (equivalent to 1500 ng fro non-split PE plasmid) together with fixed amounts of pegRNA (500 ng) and ngRNA (245 ng) plasmids were transfected. Three days post-transfection, cells were harvested, and genomic DNA was extracted using QuickExtract™ DNA Extraction Solution (Lucigen) according to the manufacturer's instructions. The genomic DNA was used for PCR amplification of the DNA flanking the prime editing target sites (see primers in Supplemental Table 2). PCR fragments were gel-purified and subjected to Sanger sequencing (Eurofins Genomics). The prime editing efficiency was analyzed by Inference of CRISPR Edits (ICE).

RNA delivery was performed by electroporation using the 4D-Nucleofector device (Lonza) in 20 μL format Nucleocuvette Strips according to the manufacturer's protocols. HEK293T cells were trypsinized just before electroporation and $2\times 10^5$ cells were used per 20 μL reaction. The RNA amounts used were: 2 ug of PE/$PE^{CO}$ mRNA, 1 ug of pegRNA and 1 ug of ngRNA. The electroporation buffer and program used for HEK293T cells were Opti-MEM (Gibco) and P3-CM138, respectively. Prime editing analysis included genomic DNA extraction and Sanger sequencing performed as described above.

**Western blot analysis**

$2.5\times 10^5$ HEK293T cells in 1 ml culture medium were seeded per well in 12-well plate one day prior to transfection. 500 ng PE plasmid was transfected using Lipofectamine 2000. Two days after transfection, the cells were lysed using the Blue Loading Buffer

Pack (#7722, Cell Signaling) according to the manufacturer's protocol. The isolated total cellular protein was electrophoresed on a 4–20% precast polyacrylamide gel (Bio-Ras) and transferred to a polyvinylidene fluoride (PVDF) membrane. PE protein was detected using an anti-Cas9 antibody (#14697, Cell Signaling) with a 1:1000 dilution according to the manufacturer's instructions. The β-actin protein was detected by β-Actin (8H10D10) Mouse mAb antibody (#3700, Cell Signaling) with a 1:1000 dilution. The signals were visualized by ChemiDoc MP Imaging System (Bio-Rad) and analyzed by Image J.

**AAV vector production**

AAV6 vectors were produced as described with a few modifications[3]. In brief, HEK293T cells were seeded at 11× $10^6$ cells per 15-cm culture dish one day before transfection. One 15-cm dish was transfected using standard PEI transfection with 6μg ITR-containing plasmid and 22μg pDGM6 (a kind gift from D. Russell), which contains the AAV6 cap genes, AAV2 rep genes, and adenovirus helper genes. AAV vectors were collected 48 hours post-transfection from cells by three freeze-thaw cycles followed by a 45 min incubation with TurboNuclease at 250U $ml^{-1}$ (Sigma). AAV vectors were purified on an iodixanol density gradient by ultracentrifugation at 237,000g for 2h at 18°C. AAV vectors were extracted at the 60-40% iodixanol interface and concentrated using a 100K Amicon Ultra -15 centrifugal filters with 1X sorbitol containing 0,001% pluronic acid. Vectors were aliquoted, and stored at -80°C until use. AAV6 vectors were titrated using Droplet Digital PCR (ddPCR; QX-200 from Bio-Rad) to quantify number of vector genomes as described previously.[4] The ITR primers and probe used for ddPCR were: Forward primer 5′-GGAACCCCTAGTGATGGAGTT-3′; Reverse primer 5′-CGGCCTCAGTGAGCGA-3′; ITR Probe: 5′-FAM-CACTCCCTCTCTGCGCGCTCG-IBFQ-3′ (IDT).

**Dual AAV infection**

10,000 HEK293T cells in 100 ul culture medium were seeded in 96-well plates and infected with two AAVs at MOIs of $5\times10^5$. Five days post-infection, cells were subjected to prime editing analysis as described above.

### References


1. Anzalone AV, *et al.* Search-and-replace genome editing without double-strand breaks or donor DNA. *Nature* **576**, 149-157 (2019).

2. Jensen TI, *et al.* Targeted regulation of transcription in primary cells using CRISPRa and CRISPRi. *Genome Research*, gr. 275607.275121 (2021).

3. Dever DP, *et al.* CRISPR/Cas9 β-globin gene targeting in human haematopoietic stem cells. *Nature* **539**, 384-389 (2016).

4. Furuta-Hanawa B, Yamaguchi T, Uchida E. Two-dimensional droplet digital pcr as a tool for titration and integrity evaluation of recombinant adeno-associated viral vectors. *Human gene therapy methods* **30**, 127-136 (2019).

5. Dang Y, *et al.* Optimizing sgRNA structure to improve CRISPR-Cas9 knockout efficiency. *Genome biology* **16**, 1-10 (2015).

**Figure legends**

**Fig S1. Screening of RT variants for prime editing activity.** The RT variants were human codon-optimized by GenScript and used to replace the M-MLV RT to form PE variants. PE constructs were transfected together with pegRNA and ngRNA plasmids into HEK293T cells and prime editing results were analyzed after 3 days. The data are presented as mean values with all data points from independent experiments shown.

**Fig S2. PE$^{CO}$, PE$^{CO}$ -ΔR and PE$^{CO}$-Mini have the same prime editing outcomes.** The representative Sanger sequencing chromatograms are from data presented in **Fig. 2B**. The control sample was not subjected to prime editing reagents and used as control for ICE analysis of Indel profiles. Note that the predominant Indels are correct prime edits and no unspecific indels were detected by ICE analysis.

**Fig S3. Schematic of the AAV vector carrying the N-terminal part of the split PE system.** The size of the AAV vector genome including the ITRs is illustrated schematically. To prevent recombination, we used two different RNA Pol III promoters (U6 and 7SK) to drive pegRNA and ngRNA expression, respectively. Also, an optimized gRNA scaffold with some sequence modifications were used for the pegRNA[5]. EFS: EF-1 Alpha Short promoter.

#### Supplemental Table 1. Spacer and 3' extension sequences of pegRNAs and spacer sequences of ngRNAs.

| | | **Spacer sequence** | **3' extension sequence** | **PBS length (nt)** | **RT template length (nt)** |
|---|---|---|---|---|---|
| pegRNA | HEK3 +1 CTT insertion | GGCCCAGACTGAGCACGTGA | AGACTTCTCCACAGGAGTCAGGTGCAC | 13 | 10 |
| | HBB +3 A to T | GCATGGTGCACCTGACTCCTG | AGACTTCTCCACAGGAGTCAGGTGCAC | 13 | 14 |
| | RNF2 +5 G to | GTCATCTTAGTCATTACCTG | AACGAACACATCAGGTAATGACTAAGATG | 15 | 14 |
| ngRNA | HEK3 +1 CTT insertion | GTCAACCAGTATCCCGGTGC | | | |
| | HBB +3 A to T | CCTTGATACCAACCTGCCCA | | | |
| | RNF2 +5 G to | TCAACCATTAAGCAAAACAT | | | |

#### Supplemental Table 2. Primers used for PCR amplification and Sanger sequencing.

| | | | |
|---|---|---|---|
| HBB +3 A to T | HBB Fw | CCAACTCCTAAGCCAGTGCCAGAAGAG | Sanger sequencing |
| | HBB Rev | AGTCAGTGCCTATCAGAAACCCAAGAG | |
| HEK3 +1 CTT insertion | HEK3 Fw | ATGTGGGCTGCCTAGAAAGG | Sanger sequencing |
| | HEK3 Rev | CCCAGCCAAACTTGTCAACC | |
| RNF2 +5 G to T | RNF2 Fw | ACGTCTCATATGCCCCTTGG | Sanger sequencing |
| | RNF2 Rev | ACGTAGGAATTTTGGTGGGACA | |

#### Supplemental Data 1: RT variants from different species

##### HIV-p51 (1320 bp)

CCTATCAGCCCCATCGAGACAGTGCCCGTGAAGCTGAAACCTGGAATGGATGGCCCTAAAGTGAAGCAATGGCCTCTGACCGAGGAAAAGATCAAGGCCCTGGTCGAGATT
TGTACAGAGATGGAAAAGGAGGGCAAGATTAGCAAGATTGGACCTGAAAACCCTTACAACACCCCTGTCTTTGCCATCAAGAAAAAAGACAGCACCAAGTGGCGGAAGCTG
GTGGACTTTAGAGAGCTGAACAAGCGGACCCAGGATTTCTGGGAGGTGCAGCTCGGCATCCCCCACCCCGCCGGCCTGAAGAAAAAAAAGAGCGTGACCGTGCTGGATGTG
GGCGACGCCTACTTCTCCGTGCCCCTGGACGAGGACTTCCGCAAGTACACCGCCTTCACAATCCCAAGCATCAACAACGAGACACCAGGAATCAGATACCAATACAATGTG
CTGCCTCAGGGCTGGAAGGGTAGCCCTGCCATCTTCCAGAGCAGCATGACCAAAATCCTGGAGCCTTTCAGAAAGCAGAACCCCGATATCGTGATCTACCAGTATATGGAC
GACCTGTACGTGGGCAGCGACCTGGAAATCGGCCAGCACAGAACAAAGATCGAGGAACTGCGGCAGCACCTCCTGAGGTGGGGACTGACTACCCCCGATAAGAAGCACCAG
AAAGAACCTCCTTTCCTGTGGATGGGCTACGAGCTGCACCCTGACAAATGGACAGTGCAGCCTATCGTGCTGCCTGAGAAGGACAGCTGGACCGTGAATGACATCCAAAAG
CTGGTTGGCAAGCTGAACTGGGCCAGCCAGATCTACCCTGGCATTAAGGTGCGGCAGCTGTGCAAGCTGCTGAGAGGGACCAAGGCGCTGACCGAAGTGATCCCTCTGACC
GAAGAGGCCGAGCTGGAACTGGCCGAGAACAGAGAAATCCTGAAGGAACCCGTGCACGGCGTGTACTACGACCCCTCTAAGGACCTGATCGCCGAGATCCAGAAGCAGGGC
CAGGGCCAGTGGACCTACCAAATCTACCAGGAGCCTTTTAAGAACCTGAAGACCGGCAAATACGCCAGAACCAGAGGCGCCCACACCAACGACGTGAAGCAGCTGACAGAA
GCCGTGCAGAAGATCACCACAGAGAGCATCGTGATCTGGGGCAAGACCCCAAAGTTCAAGCTGCCAATCCAGAAGGAAACATGGGAAACATGGTGGACCGAGTACTGGCAG
GCTACATGGATCCCTGAGTGGGAGTTCGTGAATACCCCTCCACTGGTCAAACTGTGGTACCAGCTGGAAAAGGAACCTATCGTAGGCGCTGAGACCTTC

##### HIV-p66 (1680 bp)

CCTATCAGCCCCATCGAGACAGTGCCCGTGAAGCTGAAACCTGGAATGGATGGCCCTAAAGTGAAGCAATGGCCTCTGACCGAGGAAAAGATCAAGGCCCTGGTCGAGATT
TGTACAGAGATGGAAAAGGAGGGCAAGATTAGCAAGATTGGACCTGAAAACCCTTACAACACCCCTGTCTTTGCCATCAAGAAAAAAGACAGCACCAAGTGGCGGAAGCTG
GTGGACTTTAGAGAGCTGAACAAGCGGACCCAGGATTTCTGGGAGGTGCAGCTCGGCATCCCCCACCCCGCCGGCCTGAAGAAAAAAAAGAGCGTGACCGTGCTGGATGTG
GGCGACGCCTACTTCTCCGTGCCCCTGGACGAGGACTTCCGCAAGTACACCGCCTTCACAATCCCAAGCATCAACAACGAGACACCAGGAATCAGATACCAATACAATGTG
CTGCCTCAGGGCTGGAAGGGTAGCCCTGCCATCTTCCAGAGCAGCATGACCAAAATCCTGGAGCCTTTCAGAAAGCAGAACCCCGATATCGTGATCTACCAGTATATGGAC
GACCTGTACGTGGGCAGCGACCTGGAAATCGGCCAGCACAGAACAAAGATCGAGGAACTGCGGCAGCACCTCCTGAGGTGGGGACTGACTACCCCCGATAAGAAGCACCAG
AAAGAACCTCCTTTCCTGTGGATGGGCTACGAGCTGCACCCTGACAAATGGACAGTGCAGCCTATCGTGCTGCCTGAGAAGGACAGCTGGACCGTGAATGACATCCAAAAG
CTGGTTGGCAAGCTGAACTGGGCCAGCCAGATCTACCCTGGCATTAAGGTGCGGCAGCTGTGCAAGCTGCTGAGAGGGACCAAGGCGCTGACCGAAGTGATCCCTCTGACC
GAAGAGGCCGAGCTGGAACTGGCCGAGAACAGAGAAATCCTGAAGGAACCCGTGCACGGCGTGTACTACGACCCCTCTAAGGACCTGATCGCCGAGATCCAGAAGCAGGGC
CAGGGCCAGTGGACCTACCAAATCTACCAGGAGCCTTTTAAGAACCTGAAGACCGGCAAATACGCCAGAACCAGAGGCGCCCACACCAACGACGTGAAGCAGCTGACAGAA
GCCGTGCAGAAGATCACCACAGAGAGCATCGTGATCTGGGGCAAGACCCCAAAGTTCAAGCTGCCAATCCAGAAGGAAACATGGGAAACATGGTGGACCGAGTACTGGCAG
GCTACATGGATCCCTGAGTGGGAGTTCGTGAATACCCCTCCACTGGTCAAACTGTGGTACCAGCTGGAAAAGGAACCTATCGTAGGCGCTGAGACCTTCTACGTGGACGGC
GCCGCTTCTAGAGAGACAAAACTTGGAAAGGCCGGATATGTGACCAATAGAGGCAGACAGAAAGTTGTGACCCTGACAGATACCACCAACCAGAAAACCGAGCTGCAGGCC
ATCCACCTGGCCCTCCAAGACTCTGGCCTCGAGGTGAACATCGTGACAGACTCTCAGTATGCCCTGGGCATCATCCAGGCCCAGCCTGATAAGTCCGAGAGCGAGCTGGTG
AACCAGATCATCGAACAGCTGATCAAGAAGGAAAAGGTCTACCTGGCTTGGGTCCCCGCTCATAAGGGAATTGGCGGCAACGAGCAGGTGGATAAGCTGGTGTCCGCCGGC
ATCCGGAAAGTGCTG

##### HIV-425 _MLV-RT-200 (1275bp)

CCTATCAGCCCCATCGAGACAGTGCCCGTGAAGCTGAAACCTGGAATGGATGGCCCTAAAGTGAAGCAATGGCCTCTGACCGAGGAAAAGATCAAGGCCCTGGTCGAGATT
TGTACAGAGATGGAAAAGGAGGGCAAGATTAGCAAGATTGGACCTGAAAACCCTTACAACACCCCTGTCTTTGCCATCAAGAAAAAAGACAGCACCAAGTGGCGGAAGCTG
GTGGACTTTAGAGAGCTGAACAAGCGGACCCAGGATTTCTGGGAGGTGCAGCTCGGCATCCCCCACCCCGCCGGCCTGAAGAAAAAAAAGAGCGTGACCGTGCTGGATGTG
GGCGACGCCTACTTCTCCGTGCCCCTGGACGAGGACTTCCGCAAGTACACCGCCTTCACAATCCCAAGCATCAACAACGAGACACCAGGAATCAGATACCAATACAATGTG
CTGCCTCAGGGCTGGAAGGGTAGCCCTGCCATCTTCCAGAGCAGCATGACCAAAATCCTGGAGCCTTTCAGAAAGCAGAACCCCGATATCGTGATCTACCAGTATATGGAC
GACCTGTACGTGGGCAGCGACCTGGAAATCGGCCAGCACAGAACAAAGATCGAGGAACTGCGGCAGCACCTCCTGAGGTGGGGACTGACTACCCCCGATAAGAAGCACCAG
AAAGAACCTCCTTTCCTGTGGATGGGCTACGAGCTGCACCCTGACAAATGGACAGTGCAGCCTATCGTGCTGCCTGAGAAGGACAGCTGGACCGTGAATGACATCCAAAAG
CTGGTTGGCAAGCTGAACTGGGCCAGCCAGATCTACCCTGGCATTAAGGTGCGGCAGCTGTGCAAGCTGCTGAGAGGGACCAAGGCGCTGACCGAAGTGATCCCTCTGACC
GAAGAGGCCGAGCTGGAACTGGCCGAGAACAGAGAAATCCTGAAGGAACCCGTGCACGGCGTGTACTACGACCCCTCTAAGGACCTGATCGCCGAGATCCAGAAGCAGGGC
CAGGGCCAGTGGACCTACCAAATCTACCAGGAGCCTTTTAAGAACCTGAAGACCGGCAAATACGCCAGAACCAGAGGCGCCCACACCAACGACGTGAAGCAGCTGACAGAA
GCCGTGCAGAAGATCACCACAGAGAGCATCGTGATCTGGGGCAAGACCCCAAAGTTCAAGCTGCCAATCCAGAAGGAAACATGGGAAACATGGTGGACCGAGTACTGGCAG
GCTACATGGATCCCTGAGTGGGAGTTCGTGAATACCCCTCCACTGGTCAAACTG

##### HPV RT (1827 bp); HPV RT-ΔR (1362 bp)

AATCAAGTCGGCCACCGGAAGATCAGACCCCACAATATCGCCACCGGCGACTACCCTCCAAGACCTCAGAAGCAGTACCCCATCAATCCCAAGGCCAAGCCTAGCATCCAG
ATCGTGATCGACGACCTGCTGAAGCAGGGCGTGCTGACCCCTCAGAACAGCACCATGAACACCCCTGTGTACCCCGTGCCTAAGCCTGATGGAAGATGGCGGATGGTGCTG
GACTACCGGGAAGTGAACAAGACAATCCCTCTGACCGCCGCACAGAACCAGCACTCTGCTGGAATCCTGGCCACAATCGTGCGGCAGAAGTACAAGACCACACTGGACCTG
GCCAACGGCTTCTGGGCCCATCCTATCACACCTGAGAGCTACTGGCTGACCGCATTCACATGGCAGGGCAAGCAGTACTGTTGGACCAGACTGCCTCAGGGCTTCCTGAAC
AGCCCTGCTCTGTTTACCGCCGACGTGGTGGATCTGCTGAAAGAGATCCCCAACGTCCAGGTGTACGTGGACGACATCTACCTGAGCCACGACGACCCCAAAGAACACGTG
CAGCAGCTGGAAAAGGTGTTCCAGATCCTGCTGCAGGCCGGCTACGTGGTGTCCCTGAAGAAGTCTGAGATCGGCCAGAAAACCGTGGAATTCCTGGGCTTCAACATCACC
AAAGAAGGCAGAGGCCTGACCGACACCTTCAAGACCAAGCTGCTGAACATCACCCCTCCAAAGGACCTGAAGCAGCTGCAGTCTATCCTGGGCCTGCTGAACTTCGCCCGG
AACTTCATCCCCAACTTCGCCGAACTGGTGCAGCCCCTGTACAACCTGATTGCCTCTGCCAAGGGCAAGTACATCGAGTGGAGCGAGGAAAACACAAAGCAGCTCAACATG
GTCATCGAGGCCCTGAACACCGCCAGCAACCTGGAAGAGAGACTGCCCGAGCAGCGGCTGGTCATCAAAGTGAACACAAGCCCTAGCGCCGGCTATGTGCGGTACTACAAC
GAGACAGGCAAGAAACCCATCATGTACCTGAACTACGTGTTCTCCAAGGCCGAGCTGAAGTTCTCCATGCTGGAAAAACTGCTGACCACCATGCACAAGGCCCTGATCAAG
GCCATGGATCTGGCCATGGGCCAAGAGATCCTGGTGTACAGCCCCATCGTGTCCATGACCAAGATCCAGAAAACTCCCCTGCCTGAGCGGAAGGCCCTGCCTATCAGATGG
ATCACCTGGATGACCTATCTGGAAGATCCCCGGATCCAGTTCCACTACGACAAGACCCTGCCAGAGCTGAAGCACATCCCCGACGTGTACACAAGCTCTCAGAGCCCCGTG
AAGCACCCCTCTCAGTATGAGGGCGTGTTCTACACAGACGGCAGCGCCATCAAGAGCCCCGATCCTACCAAGAGCAACAACGCCGGCATGGGAATCGTGCACGCCACCTAC
AAGCCTGAGTACCAGGTGCTGAACCAGTGGTCTATCCCTCTGGGCAACCACACAGCCCAGATGGCCGAAATTGCCGCTGTGGAATTCGCCTGCAAGAAAGCCCTGAAGATC
CCTGGACCTGTGCTCGTGATCACCGACAGCTTTTACGTGGCCGAGAGCGCCAACAAAGAGCTGCCCTACTGGAAGTCCAATGGCTTCGTGAACAACAAGAAGAAGCCCCTC
AAGCACATCAGCAAGTGGAAGTCTATCGCCGAGTGCCTGAGCATGAAGCCCGACATCACAATCCAGCACGAGAAGGGCATCAGCCTGCAGATCCCCGTGTTCATCCTGAAG
GGAAACGCCCTGGCCGACAAGCTGGCTACACAGGGCAGCTACGTCGTGAAC

##### MMTV (Mouse mammary tumor virus) RT (1725 bp); MMTV RT-ΔR (1338 bp)

ATGATCGGCGCCATCGAGAGCAACCTGTTCGCCGATCAGATCAGCTGGAAGTCCGACCAGCCTGTGTGGCTGAATCAGTGGCCCCTGAAGCAAGAGAAGCTGCAGGCCCTG
CAACAGCTGGTCACAGAACAGCTGCAACTGGGCCACCTGGAAGAGAGCAACAGCCCCTGGAATACCCCTGTGTTCGTGATCAAGAAAAAGTCCGGCAAGTGGCGGCTGCTG
CAGGATCTGAGAGCCGTGAATGCCACCATGCACGACATGGGAGCACTGCAGCCTGGACTGCCTTCTCCTGTGGCTGTGCCTAAAGGCTGGGAGATCATCATCATCGACCTG
CAGGACTGCTTCTTCAACATCAAGCTGCACCCCGAGGACTGCAAGAGATTCGCCTTCAGCGTGCCCTCTCCAAACTTCAAGCGGCCCTACCAGCGGTTCCAGTGGAAAGTT

CTGCCCCAGGGCATGAAGAACAGCCCCACACTGTGCCAGAAATTCGTGGACAAGGCCATCCTGACCGTGCGGGACAAGTACCAGGACAGCTACATCGTGCACTACATGGAC
GACATCCTGCTGGCTCACCCCAGCAGATCCATCGTGGATGAGATCCTGACCAGCATGATTCAGGCTCTGAACAAGCACGGCCTGGTGGTGTCCACCGAGAAGATCCAGAAG
TACGACAACCTGAAGTACCTGGGCACCCACATTCAGGGCGACGCCGTGTCTTACCAGAAACTGCAGATCCGGACCGACAAGCTGCGGACCCTGAACGACTTTCAGAAGCTG
CTGGGCAACATCAACTGGATCAGACCCTTCCTGAAGCTGACCACCGGCGAGCTGAAGCCCCTGTTCGAGATTCTGAACGGCGACAGCAACCCCATCAGCATCAGAAAGCTG
ACACCTGAGGCCTGCAAAGCCCTGCAGCTCGTGAACGAGAGACTGTCTATCGCCAGAGTGAAGCGGCTGGACCTGAGCAGACCTTGGAGCCTGTGCATCCTGAAAACCGAG
TACACCCCTACCGCCTGCCTGTGGCAAAATGGCGTGCTGGAATGGATTCATCTGCCCCACATCAGCCCCAAAGTGATTACCCCTTACGACATCTTCTGCACCCAGCTCATC
ATCAAGGGCAGACACAGAAGCAAAGAGCTGTTCAGCAAGGACCCCGATTACATCGTGGTGCCCTACACCAAGGTGCAGTTCGACCTGCTGCTCCAAGAGAAAGAGGACTGG
CCTATCAGCCTGCTGGGCTTCCTGGGCGAAGTGCATTTCCATCTGCCTAAGGACCCTCTGCTGACATTCACACTGCAGACCGCCATCATCTTCCCACACATGACCAGCACC
ACACCTCTGGAAAAGGGCATCGTGATCTTCACCGACGGCAGCGCCAATGGCAGAAGCGTGACCTATATCCAGGGCAGAGAGCCCATCATCAAAGAGAACACCCAGAACACC
GCTCAGCAGGCCGAAATCGTGGCCGTGATCACCGCCTTTGAGGAAGTGTCCCAGAGCTTCAACCTGTACACCGACAGCAAATACGTGACCGGACTGTTCCCCGAGATCGAG
ACAGCCACACTGAGCCCCAGGACCAAGATCTACACCGAGCTGCGGCATCTGCAGCGGCTGATCCACAAGAGACAAGAGAAGTTCTACATCGGCCACATCCGGGGCCATACA
GGACTGCCTGGACCACTGGCTCAGGGCAATGCCTATGCCGATAGCCTGACCAGAATCCTG

#### ALV(Avian leukosis virus) RT; ALV RT-ΔR (1335 bp)

ACAGTGGCTCTGCACCTGGCCATTCCTCTGAAGTGGAAGCCCGATCACACCCCTGTGTGGATCGATCAGTGGCCTCTGCCTGAGGGAAAGCTGGTGGCTCTGACACAGCTG
GTGGAAAAAGAACTGCAGCTGGGCCACATCGAGCCCAGCCTGTCTTGTTGGAACACCCCAGTGTTCGTGATCCGGAAGGCCAGCGGCAGCTATAGACTGCTGCATGACCTG
AGAGCCGTGAACGCCAAGCTGGTGCCTTTTGGAGCTGTTCAGCAGGGCGCTCCTGTGCTGTCTGCACTTCCTAGAGGATGGCCCCTGATGGTGCTGGACCTGAAGGACTGC
TTCTTCAGCATCCCTCTGGCCGAGCAGGACAGAGAGGCCTTCGCTTTTACACTGCCCAGCGTGAACAATCAGGCCCCTGCCAGACGGTTCCAGTGGAAAGTTCTGCCCCAG
GGCATGACATGCAGCCCCACAATCTGTCAGCTGGTCGTCGGACAGGTGCTGGAACCCCTGAGACTGAAGCACCCCAGCCTGAGAATGCTGCACTACATGGACGATCTGCTG
CTGGCCGCCTCTTCTCACGATGGACTTGAAGCCGCTGGCGAGGAAGTGATCAGCACCCTGGAAAGAGCCGGCTTCACAATCAGCCCCGACAAGATCCAGAGAGAGCCCGGC
GTTCAGTACCTGGGCTATAAGCTGGGCAGCACCTATGTGGCCCCTGTTGGACTGGTGGCCGAGCCTAGAATTGCCACACTGTGGGACGTGCAGAAACTCGTGGGAAGCCTG
CAGTGGCTCAGACCCGCTCTTGGAATCCCTCCTAGACTGATGGGCCCCTTCTACGAGCAGCTGAGAGGCAGCGATCCCAACGAGGCCAGAGAATGGAACCTGGACATGAAG
ATGGCCTGGCGCGAGATCGTGCAGCTGTCTACAACAGCCGCTCTGGAAAGATGGGACCCCGCTTTGCCTCTGGAAGGCGCTGTGGCTAGATGTGAACAGGGCGCCATTGGA
GTGCTCGGACAGGGCCTTTCTACACACCCTAGACCTTGCCTGTGGCTGTTCTCCACACAGCCCACCAAGGCCTTTACCGCCTGGCTGGAAGTTCTGACCCTGCTGATTACC
AAGCTGAGGGCCTCTGCTGTGCGGACCTTTGGCAAAGAGGTGGACATCCTTCTGCTGCCTGCCTGCTTCAGAGAGGATCTGCCACTGCCTGAAGGCATCCTGCTGGCCCTG
AGAGGATTCGCCGGAAAGATCAGAAGCAGCGACACCCCTAGCATCTTCGATATCGCCAGACCTCTGCACGTGTCCCTGAAAGTGCGCGTGACAGATCACCCTGTGCCTGGA
CCTACCGCCTTCACAGATGCCAGCAGCTCTACCCACAAAGGCGTGGTCGTTTGGAGAGAAGGCCCCAGATGGGAGATCAAAGAGATCGCCGATCTGGGCGCCTCTGTGCAG
CAACTGGAAGCTAGAGCAGTGGCCATGGCTCTGCTGCTCTGGCCTACCACACCTACCAACGTGGTCACCGATTCTGCCTTCGTGGCCAAGATGCTGCTGAAGATGGGCCAA
GAGGGCGTGCCATCTACAGCCGCCGCTTTCATTCTGGAAGATGCCCTGAGCCAGAGAAGCGCCATGGCTGCTGTTCTGCATGTGCGGTCCCATTCTGAGGTGCCCGGCTTT
TTCACCGAGGGCAATGATGTGGCCGATAGCCAGGCCACCTTTCAGGCCTAT

#### XMRV(Xenotropic MuLV-related virus) RT; XMRV RT-ΔR (1542 bp)

ACCCTGAACATCGAGGACGAGTACCGGCTGCACGAGACAAGCAAAGAACCCGATGTGCCTCTGGGCAGCACCTGGCTGTCTGATTTTCCACAGGCCTGGGCCGAGACAGGC
GGAATGGGACTTGCTGTTAGACAGGCCCCTCTGATCATCCCTCTGAAGGCCACAAGCACCCCTGTGTCCATCAAGCAGTACCCCATGAGCCAAGAGGCCCGGCTGGGAATC
AAGCCCCACATTCAGAGACTGCTGGACCAGGGCATCCTGGTGCCTTGTCAGAGCCCTTGGAATACCCCTCTGCTGCCCGTGAAGAAGCCCGGCACCAACGATTACAGACCC
GTGCAGGACCTGCGGGAAGTGAACAAGAGAGTGGAAGATATTCACCCCACCGTGCCGAATCCTTACAACCTGCTGTCTGGCCTGCCTCCTAGCCACCAGTGGTACACAGTG
CTGGACCTGAAGGACGCCTTCTTCTGTCTGCGGCTGCACCCTACAAGCCAGCCTCTGTTTGCCTTCGAGTGGCGGGATCCTGAGATGGGCATTAGCGGACAGCTGACCTGG
ACCAGACTGCCCCAGGGCTTCAAGAACAGCCCCACACTGTTTGACGAGGCCCTGCACAGAGATCTGGCCGACTTCAGAATTCAGCACCCCGACCTGATCCTGCTGCAGTAC
GTTGACGATCTGCTGCTGGCCGCCACAAGCGAGCAGGATTGTCAGAGAGGAACACGGGCTCTGCTGCAGACCCTGGGCAATCTGGGATATAGAGCCAGCGCCAAGAAGGCC
CAGATCTGCCAGAAACAAGTGAAGTACCTGGGCTACCTGCTGAAAGAGGGCCAGCGTTGGCTGACCGAGGCCAGAAAAGAAACCGTGATGGGCCAGCCTACACCTAAGACA
CCCAGACAGCTGAGAGAGTTCCTGGGCACCGCCGGATTCTGCAGACTGTGGATTCCTGGCTTCGCCGAGATGGCCGCTCCTCTGTATCCTCTGACCAAGACCGGCACACTG
TTCAACTGGGGCCCTGATCAGCAGAAGGCCTACCAAGAGATCAAGCAGGCCCTGCTGACAGCCCCTGCTCTGGGACTTCCTGACCTGACCAAGCCTTTCGAGCTGTTCGTG
GACGAGAAGCAGGGCTATGCCAAAGGCGTGCTGACACAGAAGCTCGGCCCTTGGAGAAGGCCAGTGGCCTACCTGAGCAAGAAACTGGATCCTGTGGCCGCTGGCTGGCCT
CCTTGTCTGAGAATGGTGGCCGCTATTGCCGTGCTGACCAAGGATGCCGGCAAGCTGACAATGGGACAGCCTCTGGTTATTCTGGCCCCTCATGCCGTGGAAGCCCTCGTG
AAACAGCCTCCTGATCGGTGGCTGAGCAACGCCAGAATGACACACTACCAGGCCATGCTGCTGGACACCGACAGAGTGCAGTTTGGCCCTGTGGTGGCCCTGAATCCTGCT
ACATTGCTGCCCCTGCCTGAGAAAGAGGCCCCTCACGATTGCCTGGAAATCCTGGCCGAAACACACGGCACCAGACCAGACCTGACCGATCAGCCTATTCCTGACGCCGAC
TACACCTGGTATACCGACGGCAGCAGCTTCCTGCAAGAAGGACAGCGTAGAGCTGGCGCCGCTGTGACCACAGAGACAGAAGTGATTTGGGCTAGAGCCCTGCCTGCCGGA
ACATCTGCTCAGAGAGCCGAGCTGATTGCCCTGACACAGGCCCTGAAAATGGCCGAGGGCAAAAAGCTGAACGTGTACACCGACTCCAGATACGCCTTCGCCACAGCTCAT
GTGCACGGCGAGATCTATCGGAGAAGAGGCCTGCTGACCAGCGAGGGCAGAGAGATTAAGAACAAGAACGAGATCCTGGCACTGCTGAAGGCCCTGTTCCTGCCTAAGCGG
CTGAGCATCATCCACTGTCCTGGCCACCAGAAGGGCAATTCCGCTGAGGCCAGAGGCAACAGAATGGCCGATCAGGCCGCTAGAGAAGCCGCCATGAAGGCCGTGCTGGAA
ACCTCTACACTGCTG

#### HLTV-1(Human T-cell leukemia virus1) RT; HLTV-1 RT-ΔR (1317 bp)

GGCCTGGAACATCTGCCTAGACCACCTGAGATCAGCCAGTTTCCTCTGAACCCCGAGAGACTGCAGGCTCTGCAGCATCTCGTTCGGAAAGCCCTGGAAGCCGGACACATC
GAGCCTTATACAGGCCCTGGCAACAACCCTGTGTTCCCCGTGAAGAAGGCCAACGGCACCTGGCGGTTTATCCACGATCTGAGAGCCACCAACAGCCTGACCGTGGATCTG
AGCAGTAGCAGCCCTGGACCTCCTGACCTCAGCAGCCTGCCTACAACACTGGCCCATCTGCAGACCATCGACCTGAAGGACGCCTTCTTTCAGATCCCTCTGCCTAAGCAG
TTCCAGCCTTACTTCGCCTTCACCGTGCCTCAGCAGTGCAATTACGGCCCAGGCACAAGATACGCCTGGAAGGTTCTGCCCCAGGGCTTCAAGAACAGCCCCACACTGTTC
GAGATGCAGCTGGCCTCTATCCTGCAGCCTATCCGGCAGGCCTTTCCTCAGTGTGTGATCCTGCAGTACATGGACGACATCCTGCTGGCTAGCCCATCTCCTGAGGATCTG
CAGCAGCTGAGCGAAGCCACAATGGCCAGCCTGATCTCTCACGGCCTGCCTGTGTCTCAGGACAAGACCCAGCAGACCCCTGGCACCATCAAGTTTCTGGGCCAGATCATC
AGCCCCAACCACATCACCTACGACGCCGTGCCTACCGTGCCAATCAGATCCAGATGGGCTCTGCCTGAACTGCAGGCCCTCCTGGGAGAGATTCAGTGGGTGTCCAAGGGC
ACCCCTACACTGAGACAGCCTCTGCACAGCCTGTACTGTGCACTGCAGGGCCACACCGATCCTCGGGACCAGATCTACCTGAATCCTAGCCAGGTGCAGTCCCTGATGCAA
CTGCAGCAGGCCCTGAGCCAGAACTGCAGATCTAGACTGGCTCAGACCCTGCCTCTGCTGGGAGCCATTATGCTGACCCTGACCGGCACCACCACCGTGGTGTTTCAGTCT
AAGCAGCAGTGGCCCCTCGTGTGGCTTCATGCTCCTCTGCCTCACACCAGCCAGTGTCCTTGGGGACAACTGCTGGCTTCTGCTGTGCTGCTGCTGGACAAGTACACCCTG
CAGAGCTATGGCCTGCTGTGCCAGACCATCCACCACAACATCAGCATCCAGACCTTCAACCAGTTTATCCAGACCAGCGATCACCCCAGCGTGCCCATTCTGCTGCACCAC
AGCCACCGGTTCAAGAACCTGGGAGCACAGACAGGCGAGCTGTGGAACACCTTCCTGAAAACCGCTGCTCCCCTGGCTCCTGTGAAGGCTCTGACCCCTGTGTTTACACTG
AGCCCCATCATCATCAACACAGCCCCTTGCCTGTTCAGCGACGGCTCTACATCTCAGGCCGCCTACATCCTGTGGGACAAGCACATCCTGTCTCAGCGGAGCTTCCCACTG
CCTCCTCCACACAAATCTGCCCAGCAGGCAGAACTGCTGGGACTGCTCCATGGACTGAGCAGCGCCAGATCTTGGCACTGCCTGAATATCTTTCTGGACAGCAAGTACCTG
TACCACTACCTGCGGACTCTGGCCCTGGGCACATTTCAGGGCAAAAGCTCTCAGGCCCCTTTCCAGGCACTGCTGCCAAGACTGCTGGCCCACAAAGTGATCTATCTGCAC
CACGTGCGGAGCCACACCAACCTGCCTGATCCTATCAGCAAGCTGAACGCCCTGACAGACGCCCTGCTGATCACCCCTATTCTGCAGCTT

#### EIAV(Equine infectious anemia virus) RT; EIAV RT-ΔR (1260 bp)

ATCGAGCTGAAAGAGGGCACAATGGGCCCTAAGATCCCTCAGTGGCCCCTGACCAAAGAGAAGCTGGAAGGCGCCAAAGAAACCGTGCAGAGACTGCTGAGCGAGGGCAAG
ATCAGCGAGGCCAGCGACAACAACCCCTACAACAGCCCCATCTTCGTGATCAAAAAGCGGAGCGGCAAGTGGCGGCTGCTGCAGGATCTGAGAGAACTGAACAAGACCGTG
CAAGTGGGCACCGAGATCTCCAGAGGACTTCCTCATCCTGGCGGCCTGATCAAGTGCAAGCACATGACCGTGCTGGACATCGGCGACGCCTACTTCACAATCCCTCTGGAC
CCTGAGTTCAGACCCTACACCGCCTTCACTATCCCCAGCATCAACCACCAAGAGCCTGACAAGAGATACGTGTGGAAGTGCCTGCCTCAGGGCTTCGTGCTGAGCCCCTAC
ATCTACCAGAAAACCCTGCAAGAGATTCTGCAGCCCTTCCGCGAGAGATACCCTGAGGTGCAGCTGTACCAGTACATGGACGACCTGTTCGTGGGCAGCAACGGCAGCAAG
AAGCAGCACAAAGAGCTGATCATCGAGCTGCGGGCCATCCTGCAGAAGGGCTTCGAGACACCCGACGACAAGCTGCAAGAGGTGCCACCTTATAGCTGGCTGGGCTACCAG
CTGTGCCCCGAGAATTGGAAGGTGCAGAAAATGCAGCTGGACATGGTCAAGAACCCCACACTGAACGACGTGCAGAAACTGATGGGCAACATCACCTGGATGAGCAGCGGA
GTGCCTGGCCTGACCGTGAAACATATCGCCGCCACCACAAAGGGCTGCCTGGAACTGAACCAGAAAGTGATCTGGACCGAGGAAGCCCAGAAAGAGCTGGAAGAGAACAAC
GAGAAGATCAAGAACGCCCAGGGCCTGCAGTACTACAACCCCGAGGAAGAGATGCTCTGCGAGGTGGAAATCACCAAGAACTACGAGGCCACCTACGTGATCAAGCAGAGC
CAGGGAATTCTGTGGGCCGGCAAGAAAATCATGAAGGCCAACAAAGGCTGGTCCACCGTGAAGAACCTGATGCTGCTGCTGCAACACGTGGCCACCGAGAGCATCACAAGA
GTGGGCAAGTGCCCCACCTTCAAGGTGCCCTTCACCAAAGAACAAGTGATGTGGGAGATGCAGAAAGGATGGTACTACAGCTGGCTGCCCGAGATCGTGTACACCCACCAG
GTGGTGCACGACGACTGGCGGATGAAGCTGGTGGAAGAACCTACCAGCGGCATCACCATCTACACCGATGGCGGCAAGCAGAACGGCGAGGGAATTGCCGCCTACGTGACC
TCCAACGGCAGGACCAAGCAGAAAAGACTGGGCCCCGTGACACATCAGGTGGCCGAAAGAATGGCCATCCAGATGGCCCTGGAAGATACCCGGGACAAGCAAGTGAACATC
GTGACCGACAGCTACTACTGCTGGAAGAACATCACCGAAGGCCTGGGCCTCGAGGGACCTCAAAATCCTTGGTGGCCCATCATCCAGAACATCCGCGAGAAAGAAATCGTG
TACTTCGCCTGGGTGCCCGGCCACAAGGGCATCTATGGAAATCAGCTGGCCGACGAGGCCGCCAAGATCAAAGAGGAAATTATGCTGGCCTACCAGGGGACGCAAATCAAA
GAAAAGCGCGACGAGGACGCCGGCTTCGACCTGTGTGTGCCCTACGATATCATGATCCCCGTGTCCGACACCAAGATCATCCCCACCGACGTGAAGATCCAGGTTCCACCT
AACAGCTTCGGCTGGGTCACAGGCAAGAGCAGCATGGCCAAACAGGGACTGCTGATCAACGGCGGCATCATCGACGAGGGCTACACCGGCGAGATCCAAGTCATCTGCACC
AACATCGGCAAGTCCAACATCAAGCTGATCGAGGGCCAGAAGTTCGCCCAGCTCATCATCCTCCAGCACCACAGCAACAGCAGACAGCCCTGGGACGAGAACAAGATCTCC
CAGAGAGGCGACAAAGGCTTCGGCAGCACTGGCGTGTTCTGGGTCGAGAATATCCAAGAGGCTCAGGACGAGCACGAGAACTGGCACACAAGCCCCAAGATCCTGGCCAGA
AACTACAAGATCCCACTGACCGTGGCCAAGCAGATCACCCAAGAA

#### AMV(Avian myeoloblastosis virus) RT; AMV RT-ΔR (1338 bp)

ACAGTGGCCCTGCACCTGGCCATCCCCCTGAAGTGGAAGCCCAACCACACCCCAGTGTGGATCGACCAGTGGCCCCTCCCCGAGGGCAAACTGGTGGCCCTGACCCAGCTG
GTGGAAAAGGAGCTGCAACTGGGCCACATCGAGCCTTCTCTGAGCTGCTGGAATACCCCTGTCTTCGTGATCAGAAAGGCCAGCGGCTCTTACAGACTGCTGCATGATCTG
AGAGCAGTTAATGCCAAGCTGGTCCCCTTCGGTGCCGTCCAGCAGGGCGCTCCTGTGCTGTCCGCCCTGCCCAGAGGCTGGCCTCTGATGGTGCTGGATCTGAAGGACTGC
TTCTTTTCTATCCCTCTGGCCGAGCAGGACAGAGAGGCCTTCGCCTTTACACTGCCTTCTGTGAACAACCAGGCTCCAGCTCGCAGATTCCAGTGGAAAGTGCTCCCCCAG
GGCATGACCTGTAGCCCCACCATCTGCCAGCTGATCGTGGGCCAGATCCTGGAACCACTGCGGCTGAAGCACCCCAGCCTGAGAATGCTGCACTACATGGACGACCTGCTG
CTTGCCGCTTCTAGCCACGACGGCCTCGAGGCCGCCGGCGAGGAAGTGATCTCCACACTGGAAAGAGCCGGATTCACCATCAGCCCTGATAAGGTGCAGCGGGAACCTGGC
GTTCAATACCTGGGCTATAAGCTGGGAAGCACGTACGTGGCCCCTGTCGGCCTCGTGGCAGAACCTAGAATCGCTACACTGTGGGACGTGCAGAAGCTTGTGGGCAGCCTG
CAGTGGCTGAGACCCGCCCTTGGAATTCCACCTAGACTGCGGGGACCTTTCTACGAGCAGCTGCGGGGCTCCGATCCTAACGAGGCCAGAGAGTGGAACCTGGACATGAAA
ATGGCCTGGCGGGAAATCGTGAGACTGAGCACAACCGCCGCCCTGGAGAGATGGGATCCTGCCCTGCCTCTGGAGGGCGCCGTGGCCCGGTGCGAACAGGGAGCTATCGGC
GTGCTGGGCCAGGGACTGAGCACCCACCCCAGACCTTGCCTGTGGCTGTTCAGCACACAGCCTACCAAGGCCTTCACCGCCTGGCTGGAAGTGCTGACACTGCTGATCACC
AAGCTGCGGGCCAGCGCCGTGCGGACCTTCGGCAAGGAAGTCGACATCCTGCTACTGCCTGCTTGTTTTAGAGATGATCTGCCACTGCCTGAGGGCATCCTGCTGGCTCTG
AGAGGCTTTGCCGGCAAGATCCGGAGCAGCGACACCCCTAGCATCTTCGATATCGCCAGACCTCTGCACGTGTCCCTGAAGGTGCGCGTGACCGACCACCCTGTGCCTGGA
CCTACCGTGTTCACCGACGCCTCATCTAGCACCCACAAGGGCGTGGTGGTGTGGAGGGAAGGCCCCCGTTGGGAGATTAAGGAAATCGCCGACCTGGGCGCTAGCGTGCAA
CAACTGGAGGCCAGAGCCGTGGCCATGGCCCTCCTGCTGTGGCCTACCACCCCTACAAACGTGGTTACAGACAGCGCCTTCGTGGCCAAGATGCTGTTGAAAATGGGCCAG
GAGGGAGTGCCTAGCACCGCCGCTGCTTTCATCCTCGAGGACGCCCTGTCTCAGAGAAGCGCTATGGCCGCCGTGCTGCATGTGCGGTCCCACAGCGAGGTGCCCGGCTTT
TTCACCGAGGGCAACGACGTGGCCGATAGCCAGGCCACATTCCAGGCCTAC

#### Gs group II intron-RT (1260 bp)

ATGGCCCTGCTGGAGCGGATCCTGGCCAGAGACAACCTGATCACCGCTCTGAAGCGGGTGGAGGCCAATCAGGGCGCCCCAGGCATCGACGGCGTCTCCACAGATCAGCTG
CGGGATTACATCCGGGCCCATTGGAGCACCATCCACGCTCAGCTGCTCGCTGGCACATACCGTCCTGCCCCCGTGCGGAGAGTTGAGATTCCTAAGCCCGGCGGAGGAACA
AGACAGCTGGGAATCCCCACCGTGGTTGATCGCCTGATCCAGCAGGCCATCCTGCAGGAGCTGACCCCTATCTTCGATCCTGACTTCAGCAGCTCTAGCTTCGGCTTCCGG
CCCGGCAGAAACGCCCACGACGCCGTGCGACAAGCCCAGGGCTATATCCAGGAGGGCTACCGGTACGTGGTGGACATGGACCTCGAGAAGTTCTTCGACAGAGTGAACCAC
GATATCCTGATGAGCAGAGTGGCCAGAAAGGTGAAGGACAAGCGAGTGCTGAAGCTGATCAGAGCCTACCTGCAAGCTGGAGTGATGATCGAGGGCGTGAAGGTGCAGACC
GAGGAAGGCACCCCTCAGGGTGGCCCTCTGAGCCCTCTGCTGGCCAACATCCTGCTGGACGACCTGGACAAGGAACTGGAAAAACGGGGCCTGAAATTTTGTAGATACGCC
GATGACTGCAACATCTACGTGAAAAGCCTGCGGGCAGGCCAGAGAGTCAAGCAGAGCATCCAGAGATTCCTGGAAAAGACCCTGAAGCTGAAAGTGAATGAGGAAAAGTCT
GCCGTGGACAGACCTTGGAAGAGAGCCTTCCTGGGCTTTTCTTTCACCCCCGAGAGAAAGGCCAGAATCCGCCTGGCCCCTCGGAGCATTCAGAGACTGAAGCAAAGAATT
AGACAGCTGACAAACCCCAACTGGTCTATCAGCATGCCTGAGCGGATCCACAGAGTGAACCAGTACGTGATGGGCTGGATCGGCTACTTTAGACTGGTGGAAACCCCTAGC
GTCCTTCAAACAATCGAGGGATGGATCAGACGGAGACTGAGACTCTGCCAGTGGCTGCAGTGGAAGAGAGTGCGGACAAGAATCAGGGAACTGAGAGCTCTGGGCCTGAAA
GAGACAGCTGTGATGGAAATCGCCAACACCAGAAAGGGCGCCTGGCGGACCACCAAGACACCACAGCTGCACCAGGCCCTGGGCAAGACCTACTGGACCGCCCAGGGACTG
AAATCCCTGACCCAGAGATATTTCGAGCTGAGACAGGGC

#### Tel4c group II intron RT (1686 bp); Tel4c group II intron RT-truncation 1 (1506 bp): Tel4c group II intron RT-truncation 1 (1209 bp)

ATGGAAACCAGGCAGATGACAGTGGATCAGACCACCGGCGCCGTGACCAACCAAACAGAGACAAGCTGGCACAGCATCAACTGGACCAAAGCCAACAGAGAGGTTAAGAGA
CTGCAGGTGCGGATCGCTAAGGCCGTTAAAGAAGGCAGATGGGGCAAGGTGAAGGCCCTGCAATGGCTGCTGACCCACTCCTTCTACGGCAAGGCTCTCGCCGTGAAACGG
GTGACCGACAACAGCGGCTCTAGAACACCCGGCGTGGACGGCATCACCTGGAGCACCCAGGAGCAGAAAACCCAAGCCATCAAAAGCCTGAGAAGAAGAGGCTACAAGCCC
CAGCCTCTGCGCAGAGTGTACATCCCTAAGGCCAACGGCAAGCAGAGACCTCTGGGCATCCCCACCATGAAGGACCGGGCTATGCAGGCCCTGTACGCCCTGGCACTTGAA
CCTGTGGCCGAGACTACAGCCGACAGAAACAGCTACGGATTCAGAAGAGGCAGATGTACCGCCGATGCCGCTGGCCAGTGCTTCCTGGCCCTGGCTAGAGCCAAGAGCGCC
GAGCACGTGCTGGACGCCGACATCTCTGGCTGTTTTGATAACATCAGCCACGAGTGGCTGCTGGCCAATACCCCTCTGGACAAGGGAATTCTGAGAAAGTGGCTGAAGTCT
GGCTTCGTGTGGAAGCAGCAGCTCTTCCCCACACACGCCGGCACCCCTCAGGGCGGCGTGATCTCCCCTGTGCTGGCCAATATCACCCTGGATGGCATGGAAGAGCTGCTG
GCTAAGCACCTGCGGGGACAGAAGGTGAACCTGATCCGGTACGCCGATGATTTCGTGGTGACCGGAAAAGATGAGGAAACCCTGGAAAAGGCCAGAAACCTGATCCAAGAG
TTCCTGAAGGAAAGAGGCCTGACACTGTCTCCTGAGAAGACAAAGATCGTGCACATCGAGGAAGGTTTTGACTTCCTGGGATGGAACATCAGAAAGTACAACGGCGTGCTG
CTGATTAAGCCTGCCAAGAAGAACGTGAAGGCCTTCCTGAAAAAGATCCGGGATACCCTGCGGGAACTGAGAACCGCCACACAGGAGATCGTGATCGACACACTCAACCCC
ATCATCAGAGGATGGGCTAACTACCACAAGGGCCAAGTCAGCAAGGAAACATTCAATAGAGTCGACTTTGCCACCTGGCATAAGCTGTGGCGGTGGGCCAGACGACGGCAT
CCTAACAAGCCAGCCCAGTGGGTGAAGGACAAGTATTTCATCAAGAACGGAAGCAGAGATTGGGTCTTTGGCATGGTGATGAAGGACAAGAATGGCGAGCTCCGGACCAAG
AGACTGATCAAGACCAGCGACACCAGAATCCAGCGGCACGTGAAGATCAAGGCCGACGCCAATCCTTTCCTCCCAGAATGGGCTGAGTACTTCGAGAAACGCAAGAAACTG
AAGAAGGCCCCAGCGCAGTATAGACGGATCCGCAGAGAGCTGTGGAAGAAACAGGGCGGCATCTGCCCTGTGTGCGGCGGAGAAATCGAGCAGGACATGCTGACAGACATC
CACCACATCCTGCCCAAGCACAAAGGCGGTTCCGACGACCTGGATAACCTGGTGCTGATCCACGCCAACTGCCACAAGCAGGTGCACAGCCGGGACGGCCAGCACTCTCGG
AGCCTGCTGAAAGAGGGCCTG

### Supplemental Data 2: codon usage M-MLV RT variants

#### IDT M-MLV RT

ACCCTTAACATAGAGGACGAATACCGACTCCACGAAACGAGCAAGGAACCTGATGTTTCTTTGGGCAGTACGTGGTTGTCCGACTTTCCGCAAGCCTGGGCGGAAACGGGG
GGTATGGGTCTCGCAGTCAGACAGGCTCCTCTTATCATTCCTTTGAAAGCGACCTCAACGCCAGTGTCAATCAAACAATACCCAATGAGTCAAGAGGCAAGGCTCGGAATA
AAGCCCCATATACAAAGGCTCCTCGATCAGGGGATCTTGGTACCATGCCAAAGCCCATGGAATACGCCCCTCCTCCCCGTCAAGAAACCGGGCACGAACGATTATAGACCG
GTGCAAGATCTCCGCGAAGTCAATAAGCGGGTGGAGGATATACACCCGACAGTACCTAACCCCTACAATCTGCTGTCTGGCCTGCCACCGTCACACCAATGGTATACCGTA
TTGGACCTGAAAGATGCCTTTTTCTGCCTGAGACTGCACCCCACCTCCCAACCATTGTTCGCATTCGAGTGGAGGGATCCAGAAATGGGGATTAGTGGCCAATTGACTTGG
ACCAGGCTCCCCCAAGGTTTTAAAAATTCACCAACTCTTTTTAATGAGGCGCTGCACAGAGACTTGGCCGACTTTAGAATTCAACACCCCGACCTCATTCTTTTGCAGTAC
GTGGATGACTTGTTGCTCGCCGCGACATCCGAACTTGATTGCCAACAGGGAACCCGCGCCCTTCTCCAAACCCTTGGTAATCTCGGCTACAGAGCCTCCGCGAAAAAGGCC
CAAATATGCCAAAAACAGGTAAAATATCTCGGTTATCTCCTGAAAGAGGGCCAGCGCTGGCTGACTGAAGCTCGCAAAGAGACCGTTATGGGTCAACCCACCCCGAAGACA
CCTAGGCAGCTGAGGGAGTTCCTGGGTAAAGCCGGATTTTGCCGGCTTTTTATACCGGGTTTTGCCGAAATGGCAGCCCCCCTCTACCCTCTGACCAAGCCAGGTACGTTG
TTTAATTGGGGACCCGATCAACAGAAAGCATATCAGGAAATTAAGCAGGCACTCCTGACTGCTCCTGCCCTTGGCCTTCCTGATCTTACGAAGCCCTTTGAGCTGTTCGTG
GACGAAAAGCAAGGGTATGCTAAAGGCGTTCTTACCCAGAAACTCGGTCCTTGGAGACGCCCGGTCGCCTACCTCTCCAAAAAGCTCGATCCTGTCGCCGCGGGCTGGCCG
CCATGTTTGCGCATGGTGGCTGCAATTGCGGTGCTGACAAAGGACGCGGGAAAGCTGACTATGGGGCAGCCGCTGGTCATCCTGGCACCGCACGCAGTTGAAGCACTCGTC
AAGCAACCTCCGGATCGCTGGCTTTCCAATGCAAGAATGACGCACTACCAAGCGCTGCTCCTTGATACTGATAGAGTACAATTTGGACCTGTCGTTGCATTGAACCCAGCG
ACTCTTCTTCCGCTGCCAGAGGAGGGCCTGCAGCATAATTGTCTGGATATCTTGGCAGAAGCTCATGGCACCCGCCCTGACTTGACCGACCAACCATTGCCTGACGCCGAC
CATACTTGGTACACTGACGGTTCTAGTCTTCTCCAGGAGGGGCAACGCAAAGCTGGCGCGGCGGTGACTACTGAAACCGAAGTGATTTGGGCTAAAGCGCTCCCTGCGGGT
ACTTCAGCACAGCGGGCAGAATTGATCGCCCTTACCCAAGCGCTTAAGATGGCCGAAGGTAAGAAGCTCAACGTTTACACGGACAGTAGGTATGCTTTTGCGACCGCACAC
ATACACGGAGAGATCTATAGACGAAGAGGGTGGCTCACATCAGAGGGTAAAGAAATTAAGAACAAAGACGAAATCTTGGCGCTCTTGAAGGCCCTTTTCCTTCCTAAAAGA
CTTAGCATTATTCATTGCCCTGGTCATCAGAAAGGACATAGCGCCGAAGCCAGAGGCAACCGCATGGCTGACCAAGCTGCCCGGAAGGCCGCAATCACTGAAACCCCAGAT
ACCTCAACCCTTCTTATCGAGAACAGCAGCCCC

#### GeneScript M-MLV RT

ACCCTGAACATCGAAGATGAGTACAGACTGCACGAGACATCTAAGGAACCTGATGTGTCCCTGGGTTCTACATGGCTGAGCGACTTCCCTCAGGCCTGGGCTGAAACCGGC
GGCATGGGCCTGGCTGTGCGGCAGGCTCCTCTGATCATCCCTCTGAAGGCCACCTCCACCCCTGTGTCCATCAAGCAGTACCCCATGAGCCAGGAGGCCAGACTGGGCATC
AAGCCCCACATCCAGCGGCTGCTCGACCAGGGAATCCTGGTCCCCTGCCAGAGCCCCTGGAACACCCCACTTTTACCTGTGAAAAAGCCCGGCACTAACGACTACCGGCCT
GTGCAGGACCTGAGAGAAGTGAACAAGCGGGTTGAGGACATCCACCCTACAGTGCCTAACCCCTACAACCTGCTGTCTGGCCTGCCACCGAGCCACCAGTGGTACACCGTG
CTGGACCTGAAAGACGCCTTCTTCTGTCTGAGACTCCATCCTACCAGCCAGCCACTGTTCGCCTTCGAGTGGCGGGACCCTGAGATGGGCATCAGCGGCCAGCTGACATGG
ACCCGGCTGCCACAGGGCTTCAAGAACAGCCCTACACTGTTCAACGAGGCCCTGCACAGAGATCTGGCTGACTTCAGAATCCAGCATCCTGACCTGATCCTGCTGCAGTAC
GTGGACGACTTGCTGCTGGCCGCTACATCTGAACTGGATTGCCAGCAAGGCACCAGAGCTCTGCTCCAGACCCTGGGCAACCTGGGGTATAGGGCCAGCGCCAAGAAGGCC
CAGATCTGCCAGAAGCAAGTGAAGTACCTGGGATACCTGCTGAAGGAAGGCCAGAGATGGCTGACCGAGGCGCGGAAGGAAACAGTGATGGGCCAGCCTACCCCTAAGACC
CCCAGACAGCTGCGAGAATTCCTGGGCAAGGCTGGCTTTTGCAGACTGTTCATCCCCGGCTTCGCTGAGATGGCCGCTCCTCTGTACCCACTGACCAAGCCCGGTACCCTG
TTCAATTGGGGCCCCGATCAGCAGAAAGCCTACCAGGAGATTAAGCAGGCCCTGCTGACCGCCCCTGCCCTCGGGCTGCCTGATCTGACAAAACCTTTCGAGCTGTTTGTC
GACGAGAAGCAGGGCTACGCCAAGGGCGTGCTGACACAGAAACTGGGACCTTGGAGAAGACCTGTGGCTTATCTCAGCAAGAAGCTGGACCCCGTGGCCGCCGGATGGCCT
CCATGTCTGAGGATGGTCGCCGCTATCGCCGTGCTGACCAAGGACGCCGGCAAACTGACCATGGGACAACCTCTGGTGATCCTGGCCCCTCACGCCGTCGAGGCCCTGGTG
AAGCAGCCTCCTGACAGATGGCTGAGCAACGCCAGAATGACCCACTACCAGGCCCTGCTGCTGGATACCGATAGAGTGCAGTTCGGCCCTGTGGTGGCTCTGAATCCTGCC
ACCCTGCTCCCACTTCCCGAGGAAGGCCTGCAGCACAACTGCCTGGACATCCTGGCAGAAGCCCACGGCACAAGACCTGACTTAACAGACCAGCCTCTGCCTGATGCCGAC
CACACCTGGTATACCGACGGCAGCAGCCTGCTGCAGGAGGGCCAAAGAAAGGCCGGAGCCGCCGTGACAACAGAAACCGAGGTGATCTGGGCCAAGGCCCTCCCTGCTGGC
ACCAGCGCCCAAAGAGCCGAGCTGATTGCCCTCACCCAGGCCCTGAAGATGGCCGAAGGCAAGAAACTGAATGTGTACACTGACTCTAGATACGCCTTTGCCACAGCCCAC
ATCCACGGCGAGATCTACAGACGGAGAGGATGGCTAACCTCTGAGGGCAAGGAAATCAAAAACAAGGATGAGATTCTGGCCCTGCTGAAGGCACTGTTTCTGCCCAAGAGA
CTGAGCATCATCCACTGCCCCGGCCACCAGAAGGGACACAGCGCCGAGGCCCGGGGCAATAGAATGGCCGATCAGGCCGCTCGGAAGGCCGCCATCACCGAAACACCCGAC
ACCAGCACACTTCTGATCGAGAACTCCAGCCCC

#### GeneArt M-MLV RT

ACCCTGAACATCGAGGACGAGTACCGGCTGCACGAGACAAGCAAAGAACCCGATGTGTCCCTGGGCAGCACCTGGCTGTCTGATTTTCCACAGGCCTGGGCCGAGACAGGC
GGAATGGGACTTGCTGTTAGACAGGCCCCTCTGATCATCCCTCTGAAGGCCACAAGCACCCCTGTGTCCATCAAGCAGTACCCCATGAGCCAAGAGGCCCGGCTGGGAATC
AAGCCCCACATTCAGAGACTGCTGGACCAGGGCATCCTGGTGCCTTGTCAGAGCCCTTGGAATACCCCTCTGCTGCCCGTGAAGAAGCCCGGCACCAACGATTACAGACCC
GTGCAGGACCTGCGGGAAGTGAACAAGAGAGTGGAAGATATTCACCCCACCGTGCCGAATCCTTACAACCTGCTGTCTGGCCTGCCTCCTAGCCACCAGTGGTACACAGTG
CTGGACCTGAAGGACGCCTTCTTCTGTCTGCGGCTGCACCCTACAAGCCAGCCTCTGTTTGCCTTCGAGTGGCGGGATCCTGAGATGGGCATTAGCGGACAGCTGACCTGG
ACCAGACTGCCCCAGGGCTTCAAGAACAGCCCCACACTGTTCAACGAGGCCCTGCACAGAGATCTGGCCGACTTCAGAATTCAGCACCCCGACCTGATCCTGCTGCAGTAC
GTTGACGATCTGCTGCTGGCCGCCACAAGCGAGCTGGATTGTCAGCAGGGAACACGCGCTCTGCTGCAGACACTGGGCAATCTGGGCTATAGAGCCAGCGCCAAGAAGGCC
CAGATCTGCCAGAAACAAGTGAAGTACCTGGGCTACCTGCTGAAAGAGGGCCAGCGTTGGCTGACCGAGGCCAGAAAAGAAACCGTGATGGGCCAGCCTACACCTAAGACA
CCCAGACAGCTGAGAGAGTTCCTGGGCAAAGCCGGATTCTGCCGGCTGTTCATCCCTGGCTTTGCCGAAATGGCCGCTCCTCTGTACCCTCTGACAAAGCCCGGAACTCTG
TTCAACTGGGGCCCAGACCAGCAGAAGGCCTACCAAGAGATTAAGCAGGCTCTGCTGACAGCCCCTGCTCTGGGACTGCCTGATCTGACCAAGCCTTTCGAGCTGTTCGTG
GACGAGAAGCAGGGCTATGCCAAAGGCGTGCTGACACAGAAGCTCGGCCCTTGGAGAAGGCCAGTGGCCTACCTGAGCAAGAAACTGGATCCTGTGGCCGCTGGCTGGCCT
CCTTGTCTGAGAATGGTGGCCGCTATTGCCGTGCTGACCAAGGATGCCGGCAAGCTGACAATGGGACAGCCTCTGGTTATTCTGGCCCCTCATGCCGTGGAAGCCCTCGTG
AAACAGCCTCCTGATCGGTGGCTGAGCAACGCCAGAATGACCCACTATCAGGCCCTGCTGCTCGACACCGACAGAGTGCAGTTTGGACCTGTGGTGGCCCTGAATCCTGCC
ACACTTCTGCCTCTGCCTGAGGAAGGCCTGCAGCACAACTGCCTGGATATCCTGGCCGAGGCTCACGGCACAAGACCCGATCTGACAGATCAGCCTCTGCCAGACGCCGAC
CACACCTGGTATACAGATGGCAGCTCCCTGCTGCAAGAAGGACAGAGAAAAGCCGGCGCTGCCGTGACCACCGAGACAGAAGTGATTTGGGCCAAAGCTCTGCCTGCCGGC
ACATCTGCTCAGAGAGCCGAACTGATTGCCCTGACACAGGCCCTGAAAATGGCCGAGGGCAAAAAGCTGAACGTGTACACCGACTCCAGATACGCCTTCGCCACCGCTCAC
ATTCACGGCGAGATCTATCGGCGGAGAGGATGGCTGACCTCTGAGGGCAAAGAGATCAAGAACAAGGACGAGATCCTGGCACTGCTGAAGGCCCTGTTCCTGCCTAAGCGG
CTGAGCATCATCCACTGTCCTGGCCACCAGAAGGGCCACTCTGCTGAGGCTAGAGGCAACAGAATGGCCGATCAGGCCGCCAGAAAGGCCGCCATTACAGAGACACCCGAC
ACCAGCACACTGCTGATCGAGAACAGCAGCCCT

#### GeneWIZ M-MLV RT

ACACTGAATATCGAGGACGAGTACAGACTGCACGAAACCAGCAAGGAGCCCGATGTGTCTCTGGGCTCCACATGGCTGAGCGATTTTCCCCAAGCTTGGGCTGAAACCGGC
GGCATGGGCCTCGCCGTGAGACAAGCCCCTCTCATCATCCCTCTGAAAGCCACCTCCACCCCCGTGAGCATCAAACAGTACCCCATGTCCCAAGAGGCCAGACTCGGCATC
AAACCCCACATCCAGAGACTCCTCGACCAAGGCATTCTGGTGCCTTGCCAGAGCCCTTGGAACACCCCTCTCCTCCCCGTGAAGAAACCCGGAACCAACGACTACAGACCC
GTGCAAGACCTCAGAGAGGTGAACAAGAGGGTCGAGGATATCCATCCCACCGTCCCTAATCCCTACAATCTGCTGAGCGGACTCCCCCCCTCCCACCAATGGTACACCGTG
CTGGACCTCAAGGACGCCTTCTTCTGTCTGAGACTCCACCCTACCTCCCAGCCTCTGTTCGCCTTCGAATGGAGAGATCCCGAGATGGGCATCTCCGGCCAGCTGACATGG
ACCAGACTCCCTCAAGGCTTCAAGAACAGCCCCACACTCTTTAACGAAGCTCTGCATAGAGATCTGGCTGATTTTAGAATCCAGCATCCCGATCTGATCCTCCTCCAATAC
GTGGACGATCTGCTGCTGGCTGCTACCAGCGAGCTCGATTGCCAGCAAGGCACCAGAGCCCTCCTCCAGACCCTCGGCAATCTCGGATATAGAGCTTCCGCTAAGAAGGCC
CAAATCTGCCAGAAGCAAGTGAAATACCTCGGCTATCTGCTGAAGGAAGGCCAGAGATGGCTGACAGAGGCTAGAAAGGAAACCGTCATGGGCCAGCCCACACCCAAAACA
CCTAGACAGCTGAGAGAATTTCTGGGCAAAGCCGGATTTTGCAGACTCTTCATCCCCGGATTCGCTGAAATGGCTGCCCCTCTCTACCCTCTGACCAAGCCCGGCACACTG

TTTAATTGGGGCCCCGACCAACAGAAGGCCTACCAAGAGATCAAGCAAGCTCTGCTGACCGCTCCCGCTCTGGGACTGCCCGACCTCACCAAGCCCTTCGAGCTGTTTGTC
GACGAAAAGCAAGGCTATGCCAAGGGCGTGCTGACCCAAAAGCTGGGCCCTTGGAGAAGACCCGTCGCCTATCTGAGCAAGAAGCTGGATCCCGTGGCCGCTGGCTGGCCT
CCTTGTCTCAGAATGGTGGCCGCTATCGCCGTGCTGACAAAAGATGCCGGAAAGCTGACCATGGGACAGCCTCTCGTCATTCTGGCTCCCCACGCCGTGGAAGCTCTGGTG
AAGCAGCCCCCCGATAGGTGGCTGTCCAACGCTAGAATGACCCACTATCAAGCTCTGCTGCTGGACACAGATAGGGTCCAGTTTGGACCCGTGGTCGCTCTGAATCCCGCC
ACACTGCTGCCTCTCCCCGAGGAGGGACTCCAGCACAACTGCCTCGACATTCTGGCCGAAGCCCACGGAACCAGACCCGATCTGACAGATCAGCCCCTCCCCGACGCCGAT
CATACATGGTACACCGATGGATCCTCTCTGCTGCAAGAGGGCCAGAGAAAGGCTGGAGCCGCCGTGACAACCGAGACAGAAGTCATCTGGGCCAAGGCTCTGCCCGCTGGA
ACCTCCGCTCAAAGAGCTGAGCTCATTGCTCTGACCCAAGCTCTGAAGATGGCCGAGGGCAAGAAGCTGAACGTCTACACCGACAGCAGATACGCCTTTGCCACCGCTCAC
ATCCACGGCGAGATCTATAGAAGGAGGGGCTGGCTCACATCCGAGGGCAAGGAGATCAAGAACAAGGACGAGATTCTGGCTCTGCTCAAAGCCCTCTTTCTGCCCAAGAGG
CTGAGCATTATCCATTGCCCCGGCCACCAGAAGGGACATTCCGCTGAGGCTAGAGGCAATAGGATGGCCGACCAAGCCGCCAGAAAAGCCGCCATCACAGAAACCCCCGAC
ACCAGCACACTGCTGATCGAAAACAGCTCCCCC

#### Twist Bioscience M-MLV RT

ACTCTTAACATCGAGGACGAATACCGCCTTCACGAAACTTCTAAGGAACCGGACGTCAGCCTTGGCTCAACCTGGCTCTCAGACTTCCCCCAAGCTTGGGCAGAGACGGGC
GGTATGGGGCTTGCTGTGAGACAGGCGCCATTGATTATTCCCTTGAAGGCTACAAGCACACCAGTAAGCATCAAGCAGTATCCGATGAGCCAGGAAGCAAGGCTTGGAATT
AAACCACATATTCAACGGCTCCTGGATCAAGGGATCCTTGTGCCTTGTCAAAGCCCATGGAATACACCACTCCTGCCTGTGAAGAAGCCGGGCACAAACGACTACAGACCC
GTACAAGACTTGCGGGAAGTGAATAAAAGGGTCGAGGATATTCATCCAACAGTACCAAATCCCTATAATCTTCTCAGTGGATTGCCCCCTTCACATCAATGGTATACCGTT
CTGGACCTTAAAGACGCGTTCTTTTGTCTCCGTTTGCATCCAACTTCTCAACCACTGTTTGCATTCGAATGGCGAGACCCCGAAATGGGTATTAGTGGTCAGCTCACATGG
ACTAGGCTTCCGCAAGGATTTAAGAATTCCCCGACACTTTTCAACGAAGCCCTGCATCGGGATCTGGCTGATTTTCGAATTCAACATCCCGATCTGATTTTGCTCCAATAT
GTTGACGATCTGTTGCTCGCTGCAACCAGCGAATTGGATTGTCAGCAGGGGACCCGTGCTCTTCTTCAGACGTTGGGCAATTTGGGATACAGAGCTTCAGCAAAGAAGGCA
CAGATCTGTCAAAAGCAAGTGAAATACCTCGGTTACCTGCTCAAGGAAGGCCAACGCTGGCTTACAGAAGCTAGGAAGGAAACCGTCATGGGACAACCGACACCTAAAACA
CCACGCCAGCTGCGCGAATTTCTCGGTAAAGCCGGATTTTGCCGGCTGTTTATACCAGGATTCGCGGAGATGGCGGCACCTCTCTATCCCCTGACAAAGCCCGGTACGCTC
TTCAACTGGGGTCCTGATCAGCAGAAAGCTTACCAGGAGATAAAACAGGCCCTCTTGACCGCGCCTGCTCTTGGCCTGCCTGACCTTACCAAACCGTTCGAGCTTTTCGTG
GATGAGAAACAAGGGTATGCAAAGGGCGTGCTGACCCAGAAGCTTGGCCCCTGGCGCAGGCCAGTAGCTTATCTCTCAAAGAAACTGGATCCCGTCGCCGCCGGATGGCCG
CCCTGTCTGCGTATGGTCGCTGCTATAGCAGTCCTTACTAAAGACGCGGGGAAATTGACAATGGGGCAACCGCTGGTTATCTTGGCACCGCACGCCGTTGAAGCTCTGGTT
AAGCAGCCCCCTGATCGGTGGCTGTCTAATGCAAGGATGACACATTACCAAGCTCTGCTCCTTGATACCGATCGTGTGCAATTTGGCCCAGTCGTGGCGTTGAATCCAGCC
ACATTGTTGCCCTTGCCAGAAGAGGGGCTCCAGCATAATTGTCTGGACATTCTCGCAGAGGCACATGGGACAAGACCTGATCTGACTGATCAACCACTGCCCGATGCAGAT
CATACTTGGTATACAGACGGCTCTTCTCTGCTGCAGGAAGGTCAAAGAAAAGCCGGCGCGGCTGTCACGACTGAAACAGAAGTGATTTGGGCAAAGGCTCTCCCTGCTGGA
ACTTCAGCACAACGCGCCGAGTTGATCGCCCTGACGCAAGCGCTGAAAATGGCCGAGGGCAAGAAATTGAACGTATACACAGACTCCAGATACGCCTTCGCCACCGCACAC
ATTCACGGTGAGATTTATCGTCGTAGAGGATGGCTTACCAGTGAGGGTAAGGAAATTAAGAACAAGGATGAAATACTGGCTCTGCTGAAGGCTTTGTTCTTGCCTAAGCGG
TTGTCCATCATACACTGCCCTGGGCACCAGAAAGGCCATTCCGCAGAAGCCCGGGGAAATAGGATGGCCGATCAGGCCGCTAGAAAAGCCGCGATTACCGAAACCCCCGAT
ACTAGTACATTGCTGATCGAGAACAGCTCTCCT

#### Benchling M-MLV RT

ACCCTTAATATTGAAGATGAGTATCGGCTACATGAAACATCAAAAGAGCCAGATGTTTCTCTTGGGTCCACATGGCTGTCTGATTTCCCTCAGGCTTGGGCGGAAACCGGG
GGCATGGGCCTGGCAGTTCGCCAGGCTCCTCTGATTATACCTCTGAAGGCAACCTCTACCCCCGTGTCCATAAAACAGTACCCCATGTCACAAGAAGCCAGACTGGGGATC
AAGCCACACATTCAGAGACTGTTGGACCAGGGAATCCTGGTACCCTGTCAGTCCCCCTGGAACACGCCCCTGCTACCCGTTAAGAAACCAGGGACTAATGATTATAGGCCT
GTTCAGGATCTGAGAGAAGTCAACAAGCGGGTGGAAGACATCCACCCCACCGTGCCCAACCCTTACAACCTCTTGAGCGGGCTCCCACCGTCCCACCAGTGGTACACTGTG
CTTGATTTAAAGGATGCCTTTTTCTGCCTGCGCCTCCACCCCACAAGTCAGCCTCTCTTCGCTTTTGAGTGGAGGGATCCAGAGATGGGAATCTCAGGACAATTGACCTGG
ACCAGACTCCCACAGGGTTTCAAAAACAGTCCTACCCTGTTTAATGAGGCACTGCACCGAGACTTAGCAGACTTCCGGATCCAGCACCCTGACTTGATCCTGTTACAGTAC
GTGGATGATTTACTGCTGGCCGCCACATCTGAGCTGGACTGCCAACAGGGTACACGGGCGCTGTTACAAACCCTAGGCAACCTCGGTTATCGGGCTTCGGCTAAGAAAGCC
CAGATTTGCCAGAAACAGGTCAAGTATCTGGGCTATCTTTTGAAAGAGGGTCAGAGATGGCTGACTGAGGCCAGAAAAGAGACTGTGATGGGGCAGCCTACTCCGAAGACC
CCTCGCCAACTCAGGGAGTTCCTGGGGAAGGCAGGCTTCTGTCGCCTCTTCATCCCTGGGTTTGCAGAAATGGCAGCCCCCCTGTACCCTCTCACCAAACCGGGCACTCTG
TTCAATTGGGGCCCAGACCAACAGAAGGCCTATCAGGAAATCAAGCAGGCTCTTTTAACTGCCCCAGCCCTGGGCTTGCCAGATTTGACTAAGCCCTTTGAGCTCTTTGTG
GACGAGAAGCAGGGCTACGCGAAAGGTGTCCTAACGCAGAAACTGGGTCCTTGGCGACGGCCGGTGGCCTACCTGAGCAAAAAGCTGGACCCAGTAGCAGCTGGGTGGCCC
CCTTGTTTGCGGATGGTCGCAGCTATTGCCGTGCTGACAAAGGATGCAGGCAAGCTTACCATGGGACAGCCACTGGTCATTCTGGCCCCCCATGCAGTGGAGGCACTAGTC
AAACAACCCCCCGACCGCTGGCTTTCCAACGCCAGGATGACTCACTATCAGGCCTTGCTTTTGGACACGGACAGGGTCCAGTTCGGACCTGTGGTAGCCCTGAATCCGGCT
ACACTGCTCCCACTGCCTGAGGAAGGGCTGCAACACAACTGCCTTGATATCCTGGCCGAAGCCCACGGAACCCGACCCGATCTTACGGACCAGCCGCTCCCAGACGCCGAC
CATACCTGGTACACGGATGGAAGCAGTCTCTTACAAGAGGGCCAGCGTAAGGCGGGAGCTGCGGTGACCACCGAAACAGAGGTGATCTGGGCTAAAGCCCTGCCAGCCGGC
ACATCCGCTCAGCGCGCTGAACTGATTGCACTCACACAGGCCCTAAAGATGGCAGAAGGTAAGAAGCTCAATGTTTACACTGATAGCCGTTATGCTTTTGCTACTGCTCAT
ATCCATGGAGAGATATACAGAAGGCGTGGGTGGCTCACATCAGAAGGCAAGGAGATTAAAAATAAAGACGAGATCTTGGCGTTGCTTAAAGCCCTCTTTCTGCCCAAAAGG
CTTAGCATTATTCATTGTCCAGGACATCAAAAGGGACACAGCGCCGAGGCTAGAGGCAACAGGATGGCTGACCAAGCGGCCCGAAAGGCAGCCATCACAGAGACTCCAGAC
ACATCTACCCTCCTCATAGAAAATAGTAGCCCC

#### Supplemental Data 3: Truncated M-MLV $RT^{CO}$

### $RT^{CO}$-ΔR (1560 bp)

```
ACCCTGAACATCGAAGATGAGTACAGACTGCACGAGACATCTAAGGAACCTGATGTGTCCCTGGGTTCTACATGGCTGAGCGACTTCCCTCAGGCCTGGGCTGAAACCGGC
GGCATGGGCCTGGCTGTGCGGCAGGCTCCTCTGATCATCCCTCTGAAGGCCACCTCCACCCCTGTGTCCATCAAGCAGTACCCCATGAGCCAGGAGGCCAGACTGGGCATC
AAGCCCCACATCCAGCGGCTGCTCGACCAGGGAATCCTGGTCCCCTGCCAGAGCCCCTGGAACACCCCACTTTTACCTGTGAAAAAGCCCGGCACTAACGACTACCGGCCT
GTGCAGGACCTGAGAGAAGTGAACAAGCGGGTTGAGGACATCCACCCTACAGTGCCTAACCCCTACAACCTGCTGTCTGGCCTGCCACCGAGCCACCAGTGGTACACCGTG
CTGGACCTGAAAGACGCCTTCTTCTGTCTGAGACTCCATCCTACCAGCCAGCCACTGTTCGCCTTCGAGTGGCGGGACCCTGAGATGGGCATCAGCGGCCAGCTGACATGG
ACCCGGCTGCCACAGGGCTTCAAGAACAGCCCTACACTGTTCAACGAGGCCCTGCACAGAGATCTGGCTGACTTCAGAATCCAGCATCCTGACCTGATCCTGCTGCAGTAC
GTGGACGACTTGCTGCTGGCCGCTACATCTGAACTGGATTGCCAGCAAGGCACCAGAGCTCTGCTCCAGACCCTGGGCAACCTGGGGTATAGGGCCAGCGCCAAGAAGGCC
CAGATCTGCCAGAAGCAAGTGAAGTACCTGGGATACCTGCTGAAGGAAGGCCAGAGATGGCTGACCGAGGCGCGGAAGGAAACAGTGATGGGCCAGCCTACCCCTAAGACC
CCCAGACAGCTGCGAGAATTCCTGGGCAAGGCTGGCTTTTGCAGACTGTTCATCCCCGGCTTCGCTGAGATGGCCGCTCCTCTGTACCCACTGACCAAGCCCGGTACCCTG
TTCAATTGGGGCCCCGATCAGCAGAAAGCCTACCAGGAGATTAAGCAGGCCCTGCTGACCGCCCCTGCCCTCGGGCTGCCTGATCTGACAAAACCTTTCGAGCTGTTTGTC
GACGAGAAGCAGGGCTACGCCAAGGGCGTGCTGACACAGAAACTGGGACCTTGGAGAAGACCTGTGGCTTATCTCAGCAAGAAGCTGGACCCCGTGGCCGCCGGATGGCCT
CCATGTCTGAGGATGGTCGCCGCTATCGCCGTGCTGACCAAGGACGCCGGCAAACTGACCATGGGACAACCTCTGGTGATCCTGGCCCCTCACGCCGTCGAGGCCCTGGTG
AAGCAGCCTCCTGACAGATGGCTGAGCAACGCCAGAATGACCCACTACCAGGCCCTGCTGCTGGATACCGATAGAGTGCAGTTCGGCCCTGTGGTGGCTCTGAATCCTGCC
ACCCTGCTCCCACTTCCCGAGGAAGGCCTGCAGCACAACTGCCTGGACATCCTGGCAGAAGCCCACGGCACAAGACCTGACTTAACAGACCAGCCTCTGCCTGATGCCGAC
CACACC
```

### $RT^{CO}$-ΔR-m (1410 bp)

```
CTGGGTTCTACATGGCTGAGCGACTTCCCTCAGGCCTGGGCTGAAACCGGCGGCATGGGCCTGGCTGTGCGGCAGGCTCCTCTGATCATCCCTCTGAAGGCCACCTCCACC
CCTGTGTCCATCAAGCAGTACCCCATGAGCCAGGAGGCCAGACTGGGCATCAAGCCCCACATCCAGCGGCTGCTCGACCAGGGAATCCTGGTCCCCTGCCAGAGCCCCTGG
AACACCCCACTTTTACCTGTGAAAAAGCCCGGCACTAACGACTACCGGCCTGTGCAGGACCTGAGAGAAGTGAACAAGCGGGTTGAGGACATCCACCCTACAGTGCCTAAC
CCCTACAACCTGCTGTCTGGCCTGCCACCGAGCCACCAGTGGTACACCGTGCTGGACCTGAAAGACGCCTTCTTCTGTCTGAGACTCCATCCTACCAGCCAGCCACTGTTC
GCCTTCGAGTGGCGGGACCCTGAGATGGGCATCAGCGGCCAGCTGACATGGACCCGGCTGCCACAGGGCTTCAAGAACAGCCCTACACTGTTCAACGAGGCCCTGCACAGA
GATCTGGCTGACTTCAGAATCCAGCATCCTGACCTGATCCTGCTGCAGTACGTGGACGACTTGCTGCTGGCCGCTACATCTGAACTGGATTGCCAGCAAGGCACCAGAGCT
CTGCTCCAGACCCTGGGCAACCTGGGGTATAGGGCCAGCGCCAAGAAGGCCCAGATCTGCCAGAAGCAAGTGAAGTACCTGGGATACCTGCTGAAGGAAGGCCAGAGATGG
CTGACCGAGGCGCGGAAGGAAACAGTGATGGGCCAGCCTACCCCTAAGACCCCCAGACAGCTGCGAGAATTCCTGGGCAAGGCTGGCTTTTGCAGACTGTTCATCCCCGGC
TTCGCTGAGATGGCCGCTCCTCTGTACCCACTGACCAAGCCCGGTACCCTGTTCAATTGGGGCCCCGATCAGCAGAAAGCCTACCAGGAGATTAAGCAGGCCCTGCTGACC
GCCCCTGCCCTCGGGCTGCCTGATCTGACAAAACCTTTCGAGCTGTTTGTCGACGAGAAGCAGGGCTACGCCAAGGGCGTGCTGACACAGAAACTGGGACCTTGGAGAAGA
CCTGTGGCTTATCTCAGCAAGAAGCTGGACCCCGTGGCCGCCGGATGGCCTCCATGTCTGAGGATGGTCGCCGCTATCGCCGTGCTGACCAAGGACGCCGGCAAACTGACC
ATGGGACAACCTCTGGTGATCCTGGCCCCTCACGCCGTCGAGGCCCTGGTGAAGCAGCCTCCTGACAGATGGCTGAGCAACGCCAGAATGACCCACTACCAGGCCCTGCTG
CTGGATACCGATAGAGTGCAGTTCGGCCCTGTGGTGGCTCTGAATCCTGCCACCCTGCTCCCACTTCCCGAGGAAGGC
```
